# Visual attention modulates the integration of goal-relevant evidence and not value

**DOI:** 10.1101/2020.04.14.031971

**Authors:** Pradyumna Sepulveda, Marius Usher, Ned Davies, Amy Benson, Pietro Ortoleva, Benedetto De Martino

## Abstract

When choosing between options, such as food items presented in plain view, people tend to choose the option they spend longer looking at. The prevailing interpretation is that visual attention increases value. However, in previous studies, ‘value’ was coupled to a behavioural goal, since subjects had to choose the item they preferred. This makes it impossible to discern if visual attention has an effect on value, or, instead, if attention modulates the information most relevant for the goal of the decision-maker. Here we present the results of two independent studies—a perceptual and a value-based task—that allow us to decouple value from goal-relevant information using specific task-framing. Combining psychophysics with computational modelling, we show that, contrary to the current interpretation, attention does *not* boost value, but instead it modulates goal-relevant information. This work provides a novel and more general mechanism by which attention interacts with choice.

## 1. Introduction

How is value constructed and what is the role played by visual attention in choice? Despite their centrality to the understanding of human decision-making, these remain unanswered questions. Attention is thought to play a central role, prioritising and enhancing which information is accessed during the decision-making process. How attention interacts with value-based choice has been investigated in psychology and neuroscience [1, 2, 3, 4, 5, 6, 7, 8, 9, 10, 11] and this question is at the core of the theory of rational inattention in economics [12, 13, 14, 15].

In this context, robust empirical evidence has shown that people tend to look for longer at the options with higher values [16, 10, 6] and that they tend to choose the option they pay more visual attention to [1, 2, 7, 3, 11]. The most common interpretation is that attention is allocated to items based on their value and that looking or attending to an option boosts its value, either by amplifying it [1, 2, 17] or by shifting it upwards by a constant amount [3]. This is formalised using models of sequential sampling to obtain the attentional drift diffusion model (aDDM), where visual attention boosts the drift rate of the stochastic accumulation processes [1]. These lines of investigation have been extremely fruitful, as they have provided an elegant algorithmic description of the interplay between attention and choice.

However, in most experiments, value is coupled to the agents’ behavioural goal: participants had to choose the item they liked the most. This makes it impossible to conclude whether the effect of attention is directly on value, as is commonly interpreted, or if attention modulates the integration of the evidence that is *most useful* for achieving a set goal. Our study aims to overcome these limitations and address the following question: is visual attention linked to value directly, or to acquiring goal-relevant information?

We designed an experimental manipulation that decouples reward value from choice by means of a simple task-framing manipulation. In the main eye-tracking part of our value-based experiment, participants were asked to choose between different pairs of snacks. We used two frame manipulations: *like* and *dislike*. In the *like* frame, they had to indicate which snack they would like to consume at the end of the experiment; this is consistent with the task used in previous studies. But in the *dislike* frame, subjects had to indicate the snack that they would prefer *not* to eat, equivalent to choosing the other option. Crucially, in the latter frame value is distinct from the behavioural goal of which item to select. In fact, in the *dislike* frame participants need to consider the “anti-value” of the item to choose the one to reject.

To anticipate our results, in the *like* frame condition we replicated the typical gaze-boosting effect: participants looked for longer at the item they were about to choose – the item they deemed most valuable. In the *dislike* frame, however, participants looked for longer at the item that they then chose to eliminate, i.e., the *least* valuable item. This means that agents paid more attention to the option they selected in the task, *not* to the option to which they deemed more valuable or wanted to consume. This suggests that attention does *not* boost value but rather is used to gather task-relevant information.

In order to understand the mechanism via which attention interacts with value in both framings, we use a dynamic accumulation model, which allows us to account for the preference formation process and its dependency on task variables (values of the options). We also show how goal-relevance shapes confidence and how confidence interacts with attention.

To test the generality of our findings we also conducted a new perceptual decision-making experiment and tested a new set of participants. In this perceptual task, participants were asked to choose between two circles filled with dots. In some blocks they had to indicate the circle with more dots – *most frame*; in others, the circle with fewer dots – *fewest frame*. In this second study we replicated all the effects of the first, value-based one, corroborating the hypothesis of a domain-general role for attention in modulating goal-relevant information that drives choice.

This work questions the dominant view in neuroeconomics about the relationship between attention and value, showing that attention does not boost value *per se* but instead modulates goal-relevant information. We conclude our work by presenting an economic model of optimal evidence accumulation. Using this model, we suggest that the behavioural strategy we observe in our experiment may be the result of deploying, in the context of binary choice, a behavioural strategy that is optimal when agents face more natural larger sets of options.

## 2. Results

In our first experiment, hungry participants (n=31) made binary choices between snacks in one of two task-frames, *like* and *dislike*. In the *like* frame, participants had to report the item they would prefer to eat; in the *dislike* frame, they chose the item they wanted to avoid eating (Figure 1A). After each choice, participants reported their confidence in having made a good choice [18, 7]. At the beginning of the experiment, participants reported the subjective value of individual items using a standard incentive-compatible BDM (see Methods).

**Figure 1.**
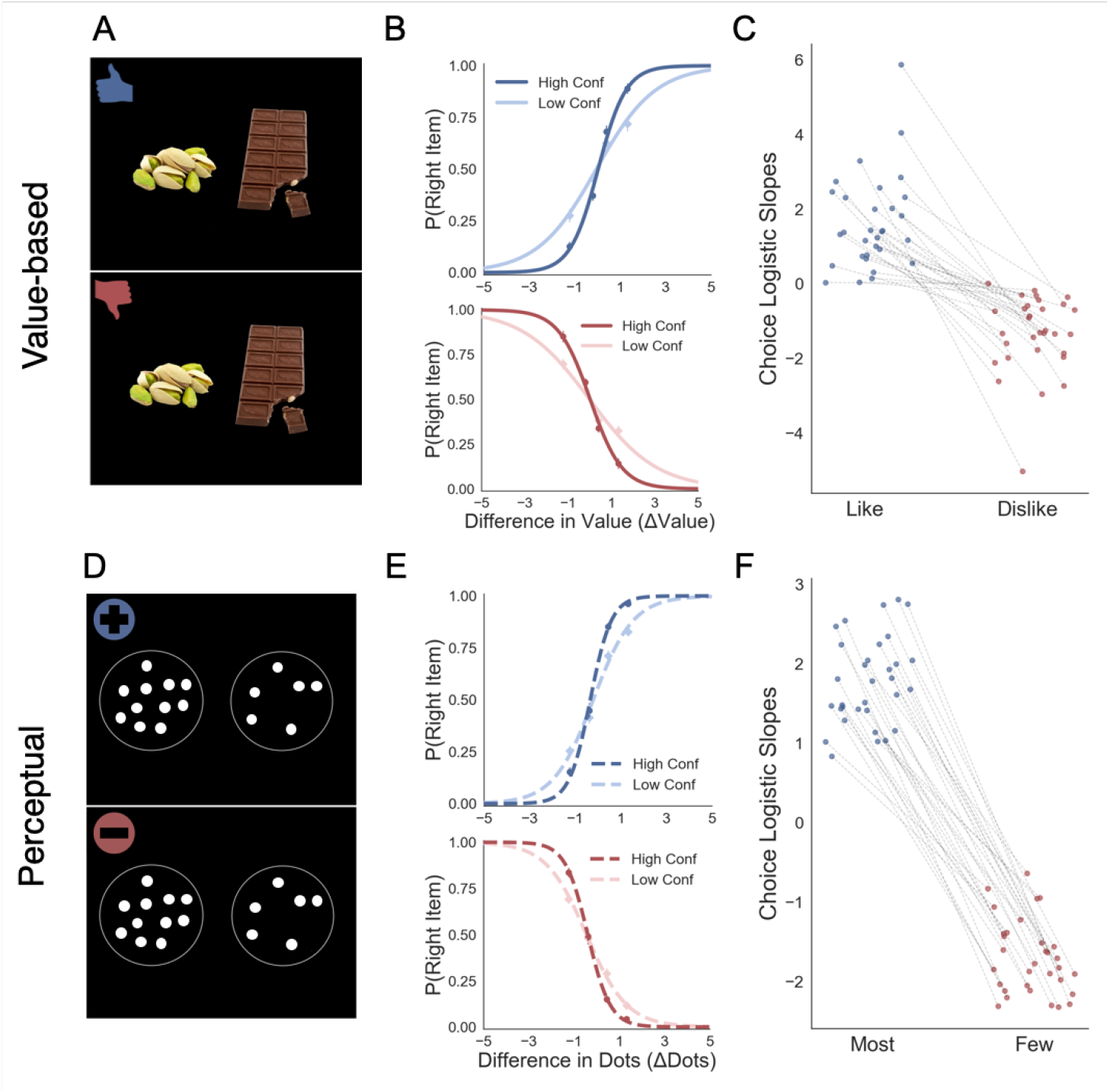
Task and behavioural results. Value-based decision task (A): participants choose between two food items presented in an eye-contingent way. Before the choice stage, participants reported the amount of money they were willing to bid to eat that snack. In the *like* frame (top) participants select the item they want to consume at the end of the experiment. In the *dislike frame* (bottom) participants choose the opposite, the item they would prefer to avoid. After each choice participants reported their level of confidence. (B) After a median split for choice confidence, a logistic regression was calculated for the probability of choosing the right-hand item depending on the difference in value (Value_Right_– Value_Left_) for *like* (top) and *dislike* (bottom) framing conditions. The logistic curve calculated from the high confidence trials is steeper, indicating an increase in accuracy. (C) Slope of logistic regressions predicting choice for each participant, depending on the frame. The shift in sign of the slope indicates that participants are correctly modifying their choices depending on the frame. Perceptual decision task (D): participants have to choose between two circles containing dots, also presented eye-contingently. In the *most* frame (top) participants select the circle with more dots. In the *fewest* frame (bottom) they choose the circle with the lower number of dots. Confidence is reported at the end of each choice. We obtained a similar pattern of results to the one observed in the Value Experiments in terms of probability of choice (E) and the flip in the slope of the choice logistic model between *most* and *fewest* frames (F).

Our second experiment was done to test whether the results observed in value-based decisions could be generalised to perceptual decisions. A different group of participants (n=32) made binary choices between two circles containing a variable number of dots (Figure 1D). In the *most* frame, participants reported the circle containing the higher number of dots; in the *fewest* frame, the one with the lower. As in the Value Experiment, at the end of each trial participants reported their confidence in their choice.

### 2.1 The effect of attention on choice

#### Value Experiment

Our results confirmed that participants understood the task and chose higher value items in the *like* frame and lower value items in the *dislike* frame (Figure 1B,C). This effect was modulated by confidence(Figure 1B) similarly to previous studies [18, 7, 19]. For a direct comparison of the differences between the goal manipulations in the two tasks (Value and Perceptual) see figure S1 and supplementary material SI 1.

We then tested how attention interacts with choice by examining the eye-tracking variables. Our frame manipulation, which orthogonalised choice and valuation, allowed us to distinguish between two competing hypotheses. The first hypothesis, currently dominant in the field, is that visual attention is always attracted to high values items and that it facilitates their choice. The alternative hypothesis is that the attention is attracted to items whose value matches the goal of the task. These two hypotheses make starkly different experimental predictions in our task. According to the first, gaze will mostly be allocated to the more valuable item independently of the frame. The second hypothesis instead predicts that in the *like* frame participants will look more at the more valuable item, while this pattern would reverse in the *dislike* frame, with attention mostly allocated to the least valuable item. In other words, according to this second hypothesis, visual attention should predict choice (and the match between value and goal) and not value, independently of the frame manipulation.

Our data strongly supported the second hypothesis since we found participants preferentially gaze (Figure 2A) the higher value option during *like* (t=7.56, *p*<0.001) and the lower value option during *dislike* frame (t=−4.99, *p*<0.001). From a hierarchical logistic regression analysis predicting choice (Figure 2B), the difference between the time participants spent observing the right over left item (ΔDT) was a positive predictor of choice both in *like* (*z*=6.448, *p*<0.001) and *dislike* (*z*=6.750, *p*<0.001) frames. This means that participants looked for longer at the item that better fit the frame and not at the item with the highest value. Notably, the magnitude of this effect was slightly lower in the *dislike* case (t=2.31, *p*<0.05). In Figure 2B are also plotted the predictors of the other variables on choice from the best fitting model.

**Figure 2.**
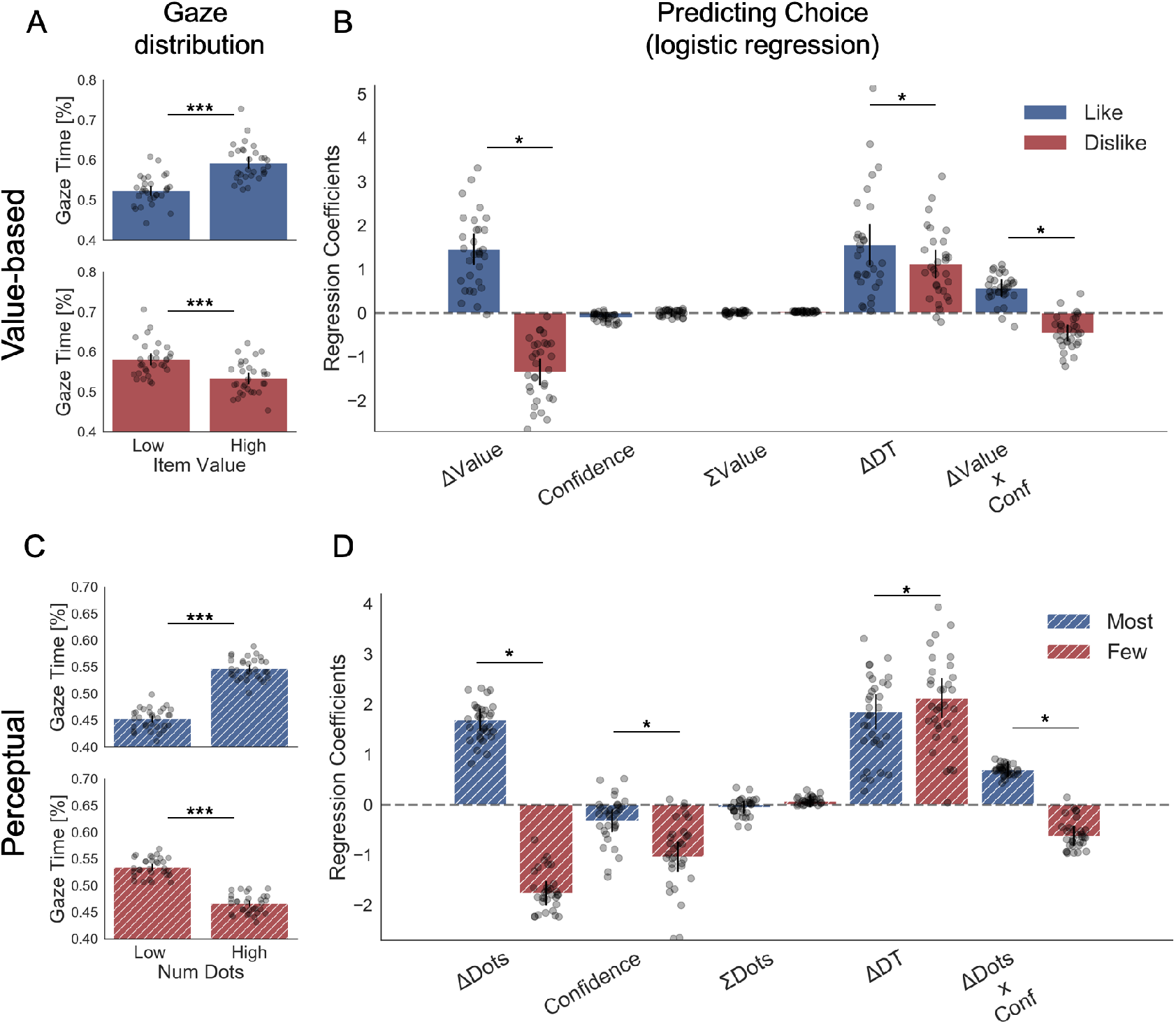
Attention and choice in Value and Perceptual Experiments. (A) Gaze allocation time depends on the frame: while visual fixations in the *like* frame go preferentially to the item with higher value (top), during the *dislike* frame participants look for longer at the item with lower value (bottom). Dots in the bar plot indicate participants’ average gaze time across trials for high and low value items. Time is expressed as the percentage of trial time spent looking at the item. Similar results were found for gaze distribution in the Perceptual Experiment (C): participants gaze the circle with higher number of dots in *most* frame and the circle with lower number of dots in *fewest* frame. Hierarchical logistic modelling of choice (probability of choosing right item) in Value (B) and Perceptual (D) Experiments, shows that participants looked for longer (ΔDT) at the item they chose in both frames. All predictors are z-scored at the participant level. In both regression plots, bars depict the fixed-effects and dots the mixed-effects of the regression. Error bars show the 95% confidence interval for the fixed effect. In Value Experiment: ΔValue: difference in value between the two items (*Value*_*Right*_ – *Value*_*Left*_); RT: reaction time; ΣValue: summed value of both items; ΔDT: difference in dwell time (DT_*Right*_– *DT*_*Left*_); Conf: confidence. In Perceptual Experiment: ΔDots: difference in dots between the two circles (*Dots*_*Right*_– *Dots*_*Left*_); ΣDots: summed number of dots between both circles. ***: *p*<0.001, **: *p*<0.01, *: *p*<0.05.

#### Perceptual Experiment

We then analysed the effect of attention on choice in the perceptual case to test the generality of our findings. As in the Value Experiment, our data confirmed that participants did not have issues in choosing the circle with more dots in the *most* frame and the one with least amount dots in the *fewest* frame (Figure 1D,F). Furthermore, as in the Value Experiment and many other previous findings [18, 7], confidence modulated the accuracy of their decisions (Figure 1E). Critically for our main hypothesis, we found that participants’ gaze was preferentially allocated to the relevant option in each frame (Figure 2C): they spent more time observing the circle with more dots during *most* frame (t=13.85, *p*<0.001) and the one with less dots during *fewest* frame (t=−10.88, *p*<0.001). ΔDT was a positive predictor of choice (Figure 2D) in *most* (z=10.249, *p*<0.001), and *fewest* (*z*=10.449, *p*<0.001) frames. Contrary to the results in the Value Experiment in which the effect of ΔDT on choice was slightly more marked in the *like* condition (Figure 2B), in the Perceptual Study the effect of ΔDT was the opposite: ΔDT had a higher effect in the *fewest* frame (ΔDT_Most-Few_: t=−2.17, *p*<0.05)(Figure 2D). However, and most importantly, in both studies ΔDT was a robust positive predictor of choice in both frame manipulations. To summarise, these results show that in the context of a simple perceptual task, visual attention also has a specific effect in modulating information processing in a goal-directed manner: subjects spend more time fixating the option they will select, not necessarily the option with the highest number of dots.

In both, Value and Perceptual Experiments, the most parsimonious models were reported in the manuscript and in Figure 2B and 2D. For a full model comparison see Figure S6 and Table S1. More details on the choice models are reported in the supplemental materials SI 2.

### 2.2 Last fixation in choice

An important prediction of attentional accumulation models is that the chosen item is generally fixated last (unless that item is much worse than the other alternative), with the magnitude of this effect related to the difference in value between the alternatives. This feature of the decision has been consistently replicated in various previous studies [1, 2, 20]. We therefore tested how the last fixation was modulated by the frame manipulation.

#### Value Experiment

In the Value Experiment in both frames we replicated the last fixation effect and its modulation by value difference between the last fixated option and the other one (Figure 3A). In the *like* frame, the probability of choosing the last item fixated upon increases when the value of the last item is higher, as is shown by the positive sign of the slope of the logistic curve (mean *β* _Like_=0.922). Crucially, during the *dislike* frame the opposite effect was found: the probability of choosing the last seen option increases when the value of the non-chosen item is higher, seen from the negative slope of the curve (mean *β* _Dislike_=−0.951; Δ*β* _Like-Dislike_: t=7.963, *p*<0.001).

**Figure 3.**
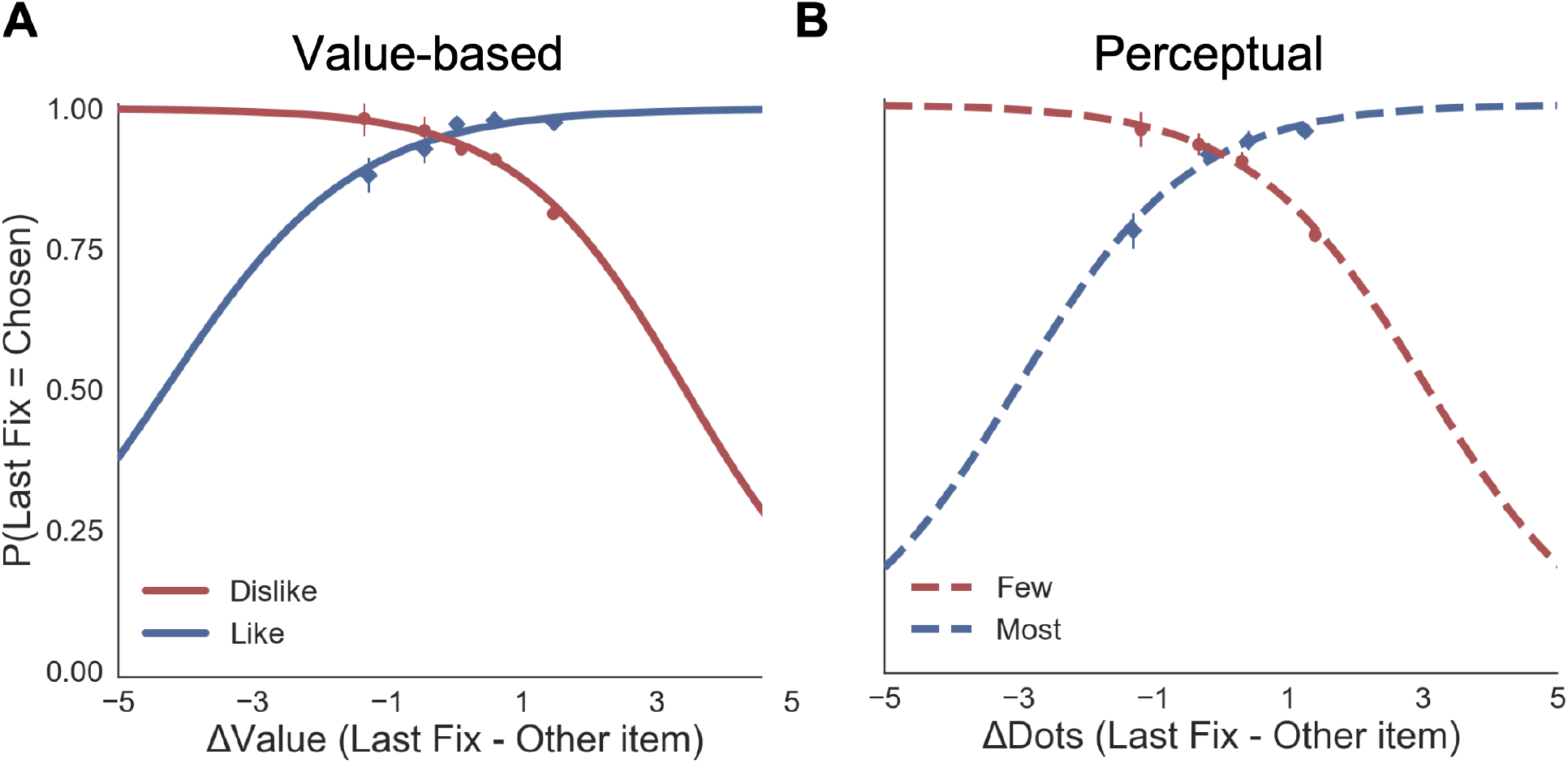
Probability that the last fixation is on the chosen item. Previous studies [1, 20] have reported a modulation of this effect by the difference in value of the items, indicating this is a prediction of evidence accumulation dynamics at the time of choice.(A) In the Value Experiment, a logistic regression was calculated for the probability the last fixation is on the chosen items (P(LastFix = Chosen)) depending on the difference in value of the item last fixated upon and the alternative item. As reported in previous studies, in *like* frame, we find it is more probable that the item last fixated upon will be chosen when the value of that item is relatively higher. In line with the hypothesis that goal-relevant evidence, and not value, is being integrated to make the decision, during the *dislike* frame the effect shows the opposite pattern: P(LastFix = Chosen) is higher when the value of the item last fixated on is lower, i.e., the item fixated on is more relevant given the frame. (B) A similar analysis in the Perceptual Experiment mirrors the results in the Value Experiment with a flip in the effect between *most* and *fewest* frames. Lines represent the model predictions and dots are the data binned across all participants. ΔValue and ΔDots measures are z-scored at the participant level.

#### Perceptual Experiment

We observed the same pattern of results that in the Value Experiment (Figure 3B). In the *most* frame, it was more probable that the last fixation was on the chosen item when the fixated circle had a higher number of dots (mean *β* _Most_=1.581). In the *fewest* frame, the effect flipped: it was more likely that the last circle seen was chosen when it had fewer dots (mean *β* _Few_ = −0.944; Δ*β* _Most-Few_: t=3.727, *p*<0.001). For further analysis of first and middle fixations see supplemental material SI 3.

### 2.3 Which factors determine confidence?

#### Value Experiment

To explore the effect that behavioural factors had over confidence, we fitted a hierarchical linear model (Figure 4A). As it was the case for the results presented above for the choice regression, the results for the confidence regression in the *like* frame replicated all the effects reported in a previous study from our lab [7]. Again, we presented here the most parsimonious model (Figure S13 and table S4 for model comparison). We found that the magnitude of ΔValue (|ΔValue|) had a positive influence on confidence in *like (z*=5.465, *p*<0.001) and *dislike* (*z*=6.300, *p*<0.001) frames, indicating that participants reported higher confidence when the items have a larger difference in value; this effect was larger in the *dislike* frame (t =−4.72, *p*<0.01). Reaction time (RT) had a negative effect on confidence in *like* (*z*=−6.373, *p*<0.001) and *dislike* (*z*=−7.739, *p*<0.001) frames, i.e., confidence was lower when the RTs were longer. Additionally, we found that, in both conditions, higher number of gaze switches (i.e., gaze shift frequence, GSF) predicted lower values of confidence in *like* (*z*=−2.365, *p*<0.05) and *dislike* (*z*=−2.589, *p*<0.05) frames, as reported in Folke et al. [7].

**Figure 4.**
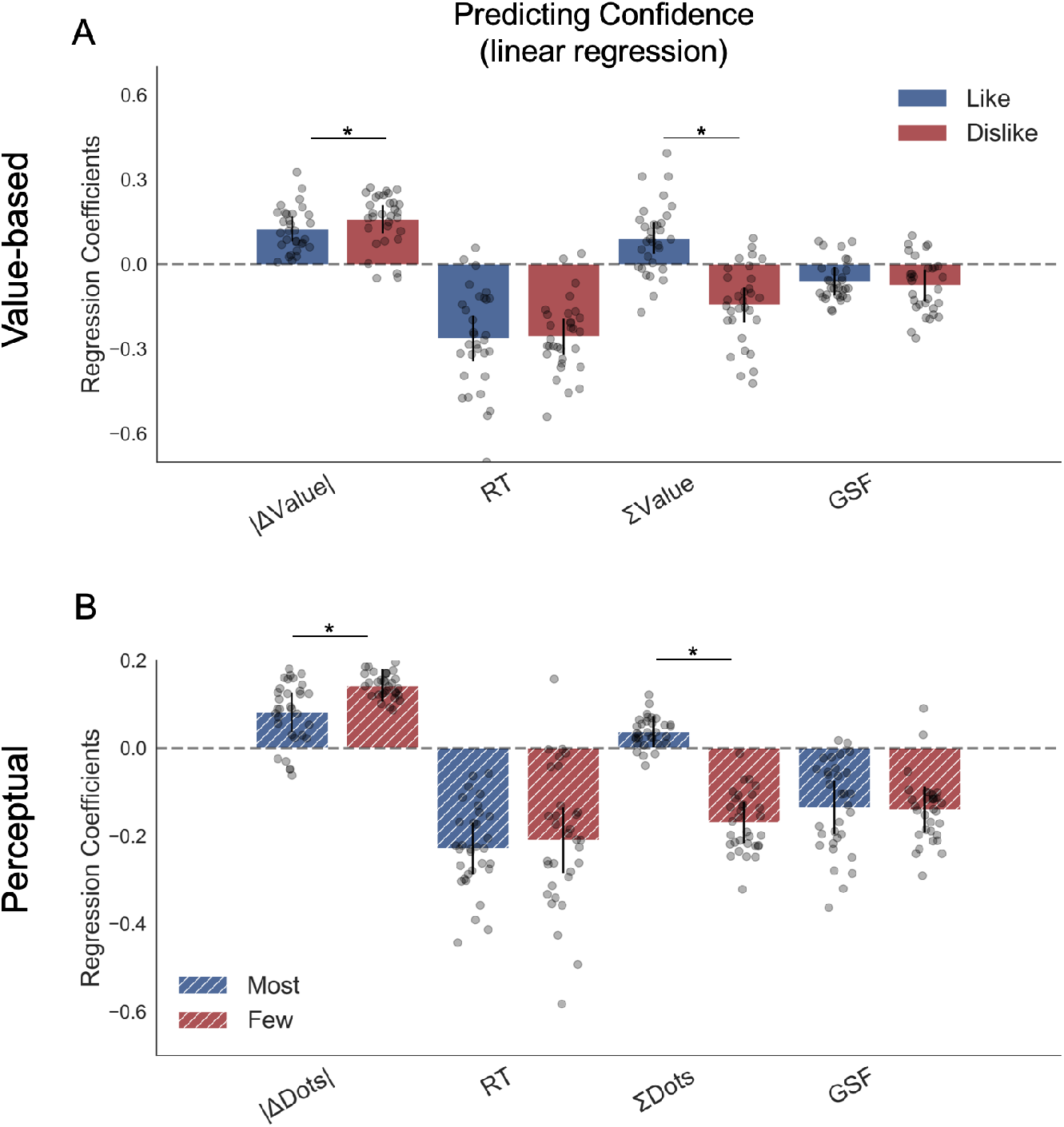
Hierarchical linear regression model to predict confidence. (A) In Value Experiment, a flip in the effect of ΣValue over confidence in the *dislike* frame was found. (B) In Perceptual Experiment a similar pattern was found in the effect of ΣDots over confidence in the *fewest* frame. The effect of the other predictors on confidence in both experiments and frames coincides with previous reports [7]. All predictors are z-scored at the participant level. In both regression plots, bars depict the fixed-effects and dots the mixed-effects of the regression. Error bars show the 95% confidence interval for the fixed effect. In Value Experiment: ΔValue: difference in value between the two items (*Value*_*Right*_– *Value*_*Left*_); RT: reaction time; ΣValue: summed value of both items; ΔDT: difference in dwell time (DT_*Right*_– *DT*_*Left*_); GSF: gaze shift frequency; ΔDT: difference in dwell time. In Perceptual Experiment: ΔDots: difference in dots between the two circles (*Dots*_*Right*_– *Dots*_*Left*_); ΣDots: summed number of dots between both circles. ***: *p*<0.001, **: *p*<0.01, *: *p*<0.05.

We then looked at the effect of the summed value of both options, ΣValue, on confidence. As in Folke et al. [7] we found a positive effect of ΣValue on confidence in the *like* frame (*z*=3.206, *p*<0.01); that is, participants reported a higher confidence level when both options were high in value. Interestingly, this effect was inverted in the *dislike* frame (*z*=−4.492, *p*<0.001), with a significant difference between the two frames (t=9.91, p<0.001) This means that, contrary to what happened in the *like* frame in which confidence was boosted when both items had high value, in the *dislike* frame confidence increased when both items had *low* value. This novel finding reveals that the change in context also generates a reassessment of the evidence used to generate the confidence reports; that is, confidence also tracks goal-relevant information.

#### Perceptual Experiment

We repeated the same regression analysis in the perceptual decision experiment, replacing value evidence input with perceptual evidence (i.e., absolute difference in the number of dots, ΔDots). We directly replicated all the results of the Value Experiment, generalising the effects we isolated to the perceptual realm (Figure 4B). Specifically, we found that ΔDots had a positive influence on confidence in *most* (*z*=3.546, *p*<0.001) and *fewest* frames (*z*=7.571, *p*<0.001), indicating that participants reported higher confidence when the evidence was stronger. The effect of absolute evidence ΔDots on confidence was bigger in the *fewest* frame (t=−4.716, *p*<0.001). RT had a negative effect over confidence in *most* (*z*=−7.599, *p*<0.001) and *fewest* frames (*z*=−5.51, *p*<0.001), i.e., faster trials were associated with higher confidence. We also found that GSF predicted lower values of confidence in *most* (*z*=−4.354, *p*<0.001) and *fewest* (*z*=−5.204, *p*<0.001) frames. Critically (like in the Value Experiment), the effect of the sum of evidence (ΣDots) on confidence also changes sign depending on the frame. While ΣDots had a positive effect over confidence in the *most* frame (z=2.061, *p*<0.05), this effect is the opposite in the *fewest* frame (*z*=−7.135, *p*<0.001), with a significant difference between the parameters in both frames (t=14.621, *p*<0.001). The magnitude of ΣDots effect was stronger in the *fewest* frame (t=−10.438, *p*<0.001). For further details on the confidence models see the supplemental information, section SI 4.

### 2.4 Attentional model: GLAM

To gain further insights into the dynamic of the information accumulation process we modelled the data from both experiments adapting a Gaze-weighted Linear Accumulator Model (GLAM) recently developed by Thomas and colleagues [11]. The GLAM belongs to the family of race models and approximates the aDDM model [1, 2] in which the dynamic aspect is discarded, favouring a more efficient estimation of the parameters. This model was chosen since, unlike the aDDM, it allowed us to test the prediction of the confidence measures as balance of evidence [21, 22, 18]. Crucially, in both experiments we used goal-relevant evidence (not the value or the number of dots) to fit the models in the *dislike* and *fewest* frames (for further details see the Methods *Attentional Model: Glam* section).

#### 2.4.1 Parameter fit and simulation

##### Value Experiment

The simulations estimated with the parameters fitted for *like* and *dislike* frames data (even-trials) reproduced the behaviour observed in the data not used to fit the model (odd-trials). In both *like* and *dislike* frames, the model replicated the observed decrease of RT when |ΔValue| is high, i.e., the increase in speed of response in easier trials (bigger value difference). The RT simulated by the models significantly correlated with the RT values observed in participants odd-numbered trials (*Like*: r=0.90, *p*<0.001; *Dislike*: r=0.89, *p*<0.001) (Figure 5A). In the *like* frame, the model also correctly predicted a higher probability of choosing the right item when ΔValue is higher. In the *dislike* frame, the model captured the change in the task goal and predicted that the selection of the right item will occur when ΔValue is higher, i.e., when the value of the left item is higher. Overall, in both frames the observed and predicted probabilities of choosing the most valuable item were significantly correlated (*Like*: r=0.80, *p*<0.001; *Dislike*: r=0.79, *p*<0.001) (Figure 5B). See figures S16A and S17A for further details.

**Figure 5.**
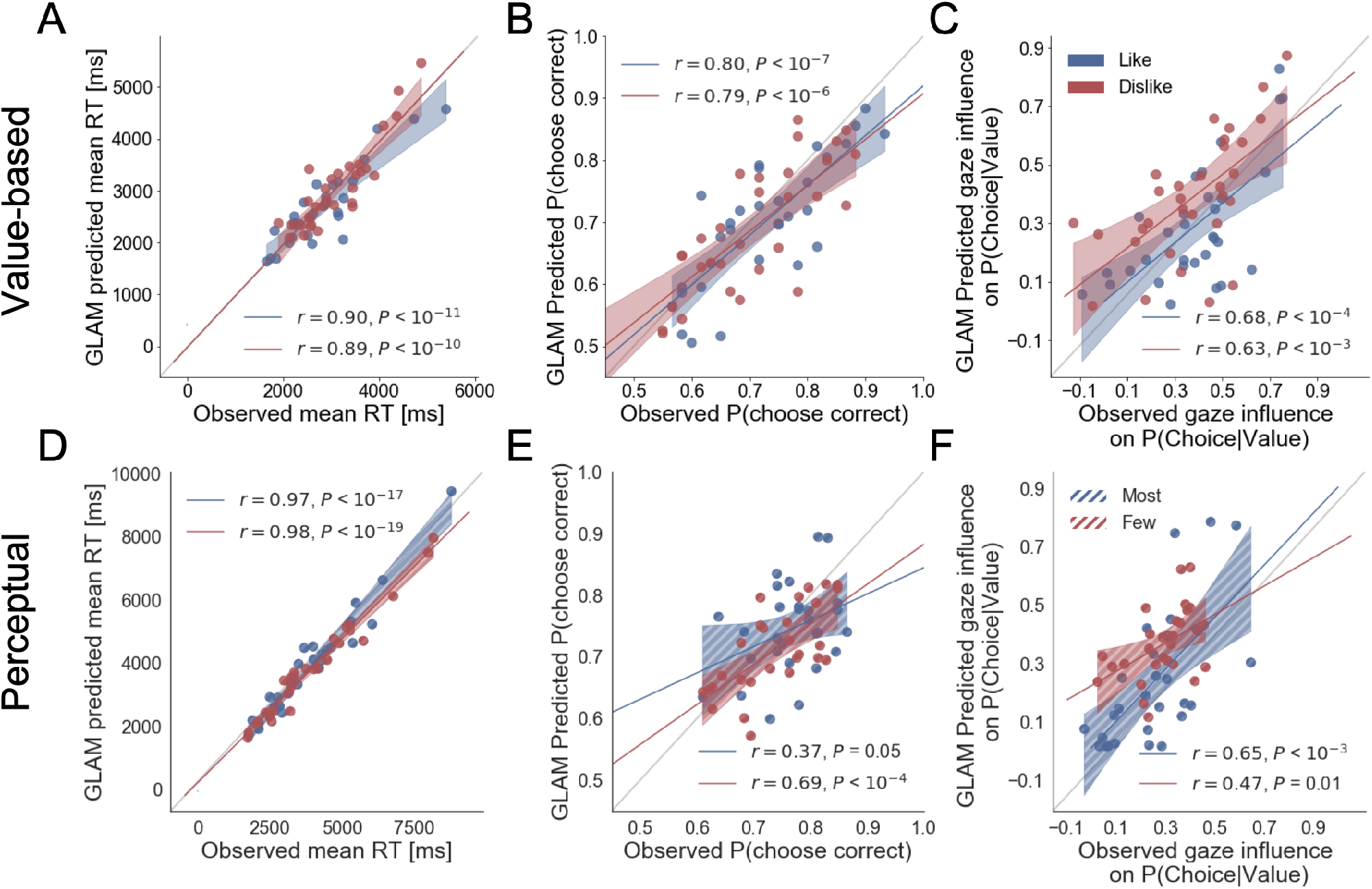
Individual out-of-sample GLAM predictions for behavioural measures in Value(A-C) and Perceptual Experiments (D-F). In value-based decision, (A) the model predicts individuals mean RT; (B) the probability of choosing the item with higher value in *like* frame, and the item with lower value in *dislike* frame; and (C) the influence of gaze in choice probability. In the Perceptual Experiment, (D) the model also predicts RT and (F) gaze influence. (E) The model significantly predicts the probability of choosing the best alternative in the *fewest* frame only (in the *most* frame a trend was found). The results corresponding to the models fitted with *like/most* frame data are presented in blue, and with *dislike/fewest* frame data in red. Dots depict the average of observed and predicted measures for each participant. Lines depict the slope of the correlation between observations and the predictions. Mean 95% confidence intervals are represented by the shadowed region in blue or red, with full colour representing Value Experiment and striped colour Perceptual Experiment. All model predictions are simulated using parameters estimated from individual fits for even-numbered trials.

In both frames, the models also predicted choice depending on the difference in gaze (ΔGaze = g_right_ - g_left_), i.e., that the probability of choosing the right item increases when the time spent observing that item is higher. However, in this case, we cannot say if gaze allocation itself is predicting choice if we do not account for the effect of |ΔValue|. To account for the relationship between choice and gaze we used a measure devised by Thomas et al. [11], ‘gaze influence’. Gaze influence is calculated taking the actual choice (1 or 0 for right or left choice, respectively) and subtracting the probability of choosing the right item given by a logistic regression for ΔValue calculated from actual behaviour. The averaged ‘residual’ choice probability indicates the existence of a positive or negative gaze advantage. Then, we compared the gaze influence predicted by GLAM with the empirical one observed for each participant. As in Thomas et al. [11], most of the participants had a positive gaze influence and it was properly predicted by the model in both frames (Like: r=0.68, *p*<0.001; *Dislike*: r=0.63, *p*<0.001) (Figure 5C).

##### Perceptual Experiment

As in the Value Experiment we fitted the GLAM to the data and we conducted model simulations. Again, these simulations showed that we could recover most of the behavioural patterns observed in participants. We replicated the relationship between RT and ΔDots (*Most*: r=0.97, *p*<0.001; *Fewest*: r=0.98, *p*<0.001) (Figure 5D). As in the value-based experiment, the model also predicted a higher probability of choosing the right-hand item when ΔDots is higher in the *most* frame and when −ΔDots is higher in the *fewest* frame. However, in the Perceptual Experiment, the simulated choices only in the *fewest* frame were significantly correlated with the observed data, although we observed a non-significant trend in the *most* frame (*Most*: r=0.69, *p*<0.001; *Fewest*: r=0.37, *p*=0.051) (Figure 5E). In both frames, we observed that the model predicted that choice was linked to ΔGaze and, as in the Value Experiment, we show that the gaze influence predicted by the model is indeed observed in the data (*Most*: r=0.65, *p*<0.001; *Fewest*: r=0.47, *p*<0.05) (Figure 5F). See figures S16B and S17B for further details.

Results of the models fitted without accounting for the change in goal-relevant evidence provided a poor fit of the data, these results are presented in the supplemental materials (Figures S14, S15 and S18). For a direct comparison of the different GLAM parameters see supplemental information SI 6. Additionally, we were able to mirror the results obtained with GLAM using aDDM [1, 8]. For *dislike* and *fewest* frames the best model was the one fitted using goal-relevant evidence (see supplemental materials SI 7 for details).

#### 2.4.2 Balance of Evidence and Confidence

The GLAM belongs to the family of race models in which evidence is independently accumulated for each option. Therefore, using the GLAM we were able to adapt the model to estimate a measure of confidence in the decision that is defined by the balance of evidence [21, 23, 22, 18] allowing us to characterise the pattern of the confidence measures. Balance of evidence is defined as the absolute difference between the accumulators for each option at the moment of choice, which is when one of them reaches the decision threshold (i.e., Δe = |E_right_(t_final_) - E_left_(t_final_)|) (Figure 6A). To estimate Δe we performed a large number of computer simulations using the fitted parameters for each participant in both experiments.

**Figure 6.**
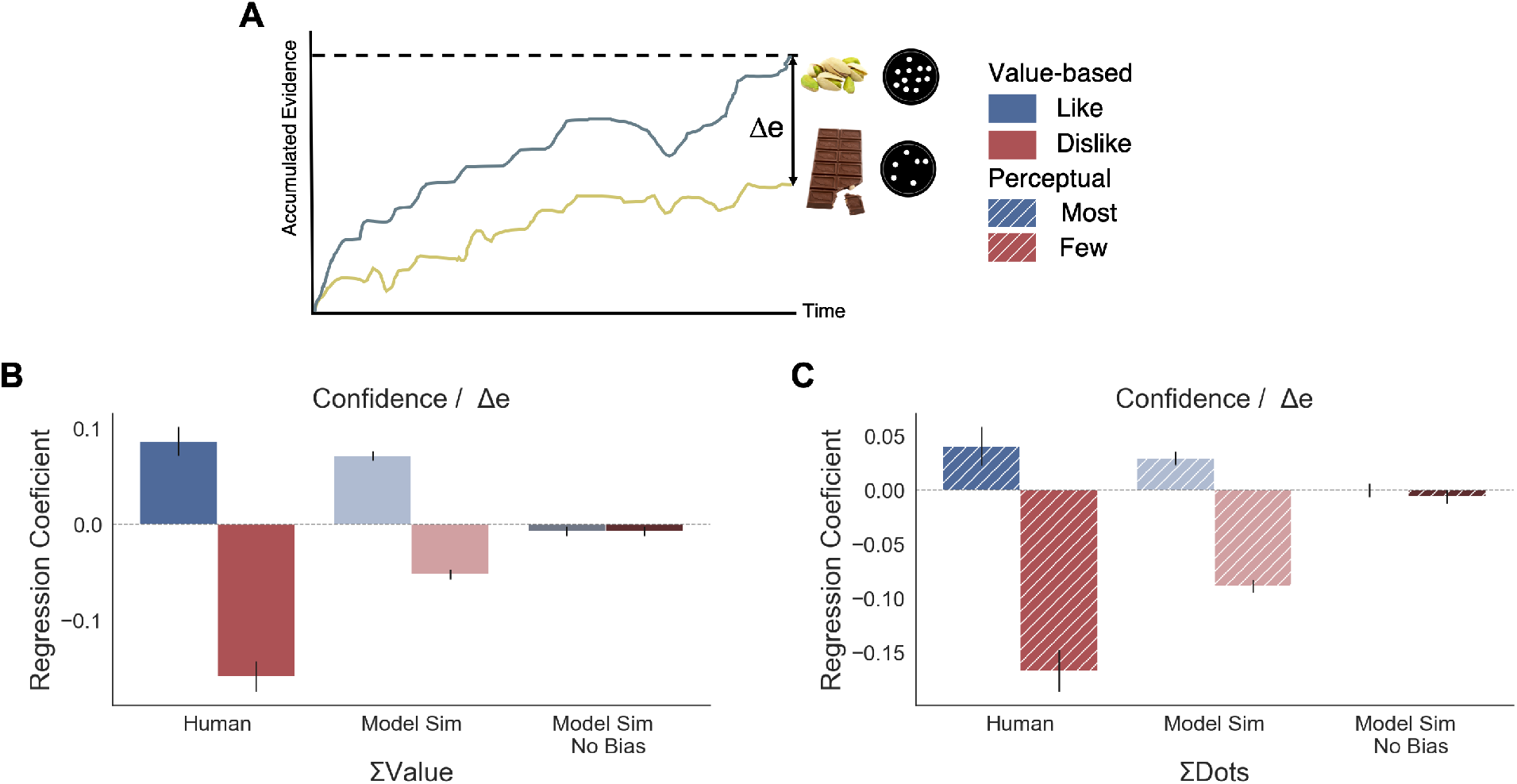
Balance of evidence (Δe) simulated with GLAM reproduces ΣValue and ΣDots effects over confidence. (A) GLAM is a linear stochastic race model in which two alternatives accumulate evidence until a threshold is reached by one of them. Δe has been proposed as a proxy for confidence and it captures the difference in evidence available in both accumulators once the choice for that trial has been made. (B) Using Δe simulations we captured the flip of the effect of ΣValue over confidence between *like* and *dislike* frames. Δe simulations were calculated using the model with parameters fitted for each individual participant. A pooled linear regression model was estimated to predict Δe. The effects of ΣValue predicting Δe are presented labelled as ‘Model Sim’. A second set of simulations was generated using a model in which no asymmetries in gaze allocation were considered (i.e., no attentional biases). This second model was not capable of recovering ΣValue effect on Δe and is labelled as ‘Model Sim No Bias’. ΣValue coefficients for a similar model using participants’ data predicting confidence are also presented labelled as ‘Human’ for comparison. (C) A similar pattern of results is found in the Perceptual Experiment, with the model including gaze bias being capable of recovering ΣDots effect on Δe. This novel effect may suggests that goal-relevant information is also influencing the generation of second-order processes, as confidence. This effect may be originated by the attentional modulation of the accumulation dynamics. Coloured bars show the parameter values for ΣValue and ΣDots and the error bars depict the standard error. Solid colour indicates the Value Experiment and striped colours indicate the Perceptual Experiment. All predictors are z-scored at participants level.

##### Value Experiment

To confirm that the relationship between confidence and other experimental variables was captured by the balance of evidence simulations, we constructed a linear regression model predicting Δe as function of the values and the RTs obtained in the simulations (Δe ~ |ΔValue| + simulated RT + ΣValue). We found that this model replicated the pattern of results we obtained experimentally (Figure 4). We then explored whether the model was able to recover the effect of ΣValue on confidence (Figure 6B). As we have shown when analysing confidence, ΣValue boosted Δe in the *like* frame (*β* _ΣValue_=0.071, t=14.21, *p*<0.001) and reduced Δe in the *dislike* frame (*β* _ΣValue_=−0.061, t=−12.07, *p*<0.001). The effect of ΣValue over confidence was replicated in the simulations with an increase of Δe when high value options are available to choose (Figure S24 and S26A,D for more details). In the *dislike* frame the fitted model also replicated this pattern of behaviour, including the adaptation to context which predicts higher Δe when both alternatives have low value. Interestingly, the replication of the effect for ΣValue over Δe with GLAM did not hold when the gaze bias was taken out of the model in *like* (*β* _ΣValue_=−0.007, t=−1.495, *p*=0.13, ns) and *dislike* (*β* _ΣValue_=−0.002, t=−0.413, *p*=0.679, ns) frames (Figure 6B). We also found that the effect of |ΔValue| on confidence was replicated by the simulated balance of evidence, increasing Δe when the difference between item values is higher (i.e., participants and the model simulations are more “confident” when |ΔValue| is higher) (Figure S24).

##### Perceptual Experiment

We conducted a set of similar analyses and model simulations in the Value Experiment (Figure 6C). We found that ΣDots boosted Δe in the *most* frame (*Most* : *β* _ΣDots_=0.029, t=4.71, *p*<0.001) and reduces Δe in the *fewest* frame (*Fewest* : *β* _ΣDots_=−0.088, t=−14.41, *p*<0.001). As in the Value Experiment this effect disappeared when the gaze bias was taken out of the model (*Most*: *β* _ΣDots_=−0.0002, t=−0.04, *p*=0.96, ns; *Fewest*: *β* _ΣDots_=−0.006, t=−1.03, *p*=0.29, ns) (see figures S25 and S26B,E for more details).

Overall, these results show how the model is capable of capturing the novel empirical effect on confidence we identified experimentally, giving computational support to the hypothesis that goal-relevant evidence is fed to second order processes like confidence. It also hints at a potential origin to the effects of the sum of evidence (i.e., ΣValue, ΣDots) on confidence, suggesting that they are caused by the asymmetries in the accumulation process generated by visual attention.

### 2.5 A Model of Optimal Information Acquisition

We then sought to understand why participants systematically accumulated evidence depending on the task at hand, instead of first integrating evidence using a task-independent strategy and then emitting a response appropriate with the task. We reasoned that this may reflect a response in line with models of rational information acquisition popular in economics. These include models of so-called rational inattention, according to which agents are rationally choosing which information to acquire considering the task, the incentives, and the cost of acquiring and processing information [12, 13, 14, 15]. As opposed to DDM or GLAM, these models attempt to investigate not only what the consequences of information acquisition are, but also *which* information is acquired.

In this model, we consider an agent facing *n* available options. Each item *i* has value *v*_*i*_ to the agent, which is unknown, and agents have a prior such that values follow an independent, identical distribution; for simplicity, we assume it to be a Normal 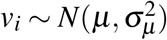. Agents can acquire information in the form of signals *x*_*i*_ = *v*_*i*_ + *ε*_*i*_, with *ε*_*i*_ independently and identically distributed with 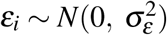. The follow Bayes’ rule in updating their beliefs after information. Once they finish acquiring information, they then choose the item with the highest expected value.

Consider first the case in which an agent needs to pick the best item out of *n* possible ones. Suppose that she already received one signal for each item. Denote *i*_1_ the item for which the agent received the highest signal, which is also the item with the highest expected value; *i*_2_ the second highest, *etc*. (Because each of these is almost surely unique, let us for simplicity assume they are indeed unique.) The agent can acquire one additional signal about any of the available items or select any probability distribution over signals. The following proposition shows that it is (weakly) optimal for the agent to acquire a second signal about the item that is currently best, i.e., *i*_1_.

Denote Δ the set of all probability distributions over signals and *V* (*i*) the utility after acquiring a new signal *x*_*i*,2_ about item *i*, i.e.,

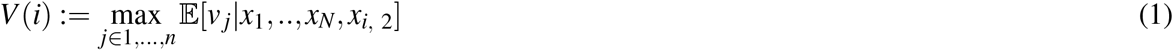

#### 1. Proposition

The optimal strategy when choosing the best option is to acquire one more signal about item i_1_ or i_2_, i.e., either the item with the currently highest expected value or the one with second highest value. That is:

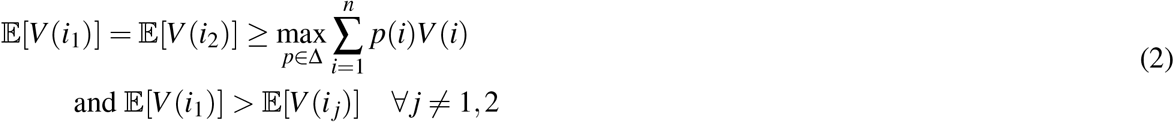

This proposition shows that agents have *asymmetric* optimal sampling strategies: they are not indifferent between which item to sample, but rather want to acquire extra signals about items that current look best or second-best. (They are indifferent between the latter two.). When *n* > 2, these strategies are strictly better than acquiring signals about any other item.

How would this change if agents need instead to pick which item to eliminate, assuming that she gets the average utility of the items she keeps? In this case, the expected utility after acquiring a new signal *x*_*i*,2_ about item *i*, is:

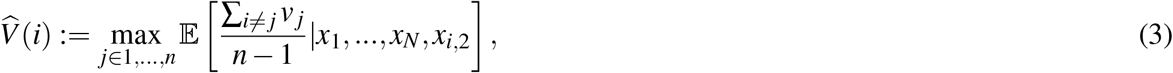

Then, it is optimal to receive an additional signal about the *least* valuable item *i*_*n*_ or the next one, *i*_*n*−1_.

#### 2. Proposition

The optimal strategy when choosing which item to discard is to acquire one more signal about item i_n_ or i_n−1_, i.e., either the one with the lowest or the one with the second lowest value. That is:

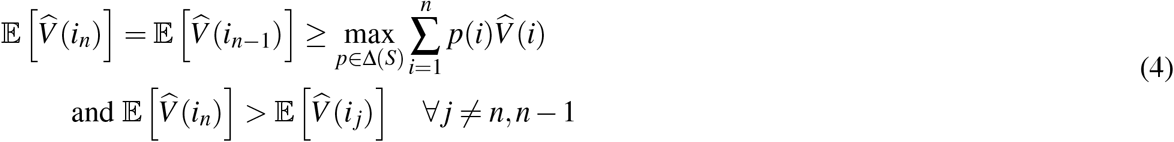

For a full proof of both propositions see supplemental materials SI 9.

Again, agents have *asymmetric* optimal sampling strategies: but now, they want to sample the items that currently look *worse* again. The intuition behind both results is that when one has to choose the best item, it is more useful to acquire information that is likely to change the ranking at the top (i.e., between best or second best item) than information that changes the ranking at the bottom, since these items won’t be selected (e.g., 4^th^ and 5^th^ item). Crucially, the reverse is true when one is tasked to select which item to eliminate.

This shows how in these simple tasks it is strictly more advantageous to acquire information in line with the current goal rather than adopting a goal-independent information-acquisition strategy.

Our model suggests that in many ecological settings, in which there are more than two options, the optimal strategy involves acquiring *asymmetric* information depending on the goal. It is only when there are only two options that individuals are indifferent about which information to acquire. We propose that the asymmetric strategies we observe even in this latter case might be a consequence of the fact that individuals have developed a strategy that is optimal for the more frequent, real-life cases in which *n* > 2, and continue to implement this same asymmetric strategy to binary choices, where it remains optimal.

## 3. Discussion

In this study we investigated how framing affects the way in which information is acquired and integrated during value-based and perceptual choices. Here, using psychophysics together with computational and economic models we have been able to adjudicate between two contrasting hypotheses. The first one, currently the dominant one in the field of neuroeconomics, proposes that attention modulates (either by biasing or boosting) a value integration that starts at the beginning of the deliberation process. Subsequently, at the time of the decision, the participant would give the appropriate response (in our task accepting the option with the highest value or rejecting the one with lowest one) using the value estimate constructed during this deliberation phase. The second hypothesis suggests that, from the very start of the deliberation process, the task-frame (goal) influences the type of information that is integrated. In this second scenario, attention is not automatically attracted to high value items to facilitate their accumulation but has a more general role in prioritising the type of information that is useful for achieving the current behavioural goal. Importantly, these two hypotheses make very distinct predictions about the pattern of attention and suggest very different cognitive architecture underpinning the decision process.

Our results favour the second hypothesis: specifically, we show that, in both perceptual and value-based tasks, attention is allocated depending on the behavioural goal of the task. While our study does not directly contradict previous findings [1, 2, 3, 17] it adds nuance to the view that this is a process specifically tied to value integration (defined as a hedonic or reward attribute). Our findings speak in favour of a more general role played by attention in prioritising whatever information is needed to fulfil a behavioural goal in both value and perceptual choices. Pavlovian influences have been proposed to play a key role in the context of accept /reject framing manipulation [24, 25, 26, 27]. However, the fact that we found almost identical results in a follow-up perceptual study in which the choice was not framed in terms of ‘accept’ or ‘reject’ but using a different kind of instruction (i.e., “choose the option with fewer or more dots”) suggests that attention acts on a more fundamental mechanism of information processing that goes beyond simple Pavlovian influences.

We also measured the trial-by-trial fluctuations in confidence to gain a deeper insight in the dynamics of this process. We found that the role of confidence goes beyond that of simply tracking the probability of an action being correct, as proposed in standard signal detection theory. Instead, it is also influenced by the perceived sense of uncertainty in the evaluation process [28, 29], and contextual cues [30]. In turn, confidence influences future responses and information seeking [7, 31, 32, 33]. In previous work [7], we reported how, in value-based choice, confidence was related not only to the difference in value between the two items, but also to the summed value (ΔValue and ΣValue using the current notation), and we found that that confidence was higher if both items have a high value [7]. Here we replicate this effect in both experiments in the *like* and *most* conditions. However, this effect flips in the *dislike* or *fewest* frame: in these cases, confidence increases when the summed value or number of dots is *smaller*. This result is particularly striking since the frame manipulation should be irrelevant for the purpose of the decision and has little effect on the objective performance. This suggests that similarly to attention, the sense of confidence is also shaped by the behavioural goal that participants are set to achieve.

In both experiments, the incorporation of goal-relevant evidence to fit the GLAM resulted in a better model fit compared with the model in which the value or perceptual evidence was integrated independently of the frame. We then modified the GLAM to include a measure of confidence defined as balance of evidence (Δe) [21, 22, 18]. In doing so we confirm that our model can replicate all the main relations between confidence, choice and RT. We then tested if the model simulation was also recovering the flip in the relationship between confidence and summed evidence (ΣValue or ΣDots) triggered by the frame manipulation. These data speak in favour of a coding scheme in which the goal sets, from the beginning of the task, the allocation of attention and, by doing so, influences first order processes such as choice, but also second order process such as confidence. Further empirical data will be required to test this idea more stringently.

The idea that the goal of the task plays a central role in shaping value-based decisions should not be surprising. Indeed, value-based decision is often called goal-directed choice. Nevertheless, there has been a surprisingly little amount of experimental work in which the behavioural goal has been directly manipulated as the key experimental variable for studying the relation between attention and value. Notable exceptions are two recent studies from Frömer and colleagues [34] and Kovach and colleagues [35] (see also supplemental analysis, Figure S7).

To gain a deeper insight into our findings we developed a normative model of optimal information acquisition rooted in economic decision theory. Our model shows that in many real-life scenarios in which the decision set is larger than two, the optimal strategy to gather and integrate information depends on the behavioural goal. Intuitively, this happens because new information all the more useful the more likely it is to change the behavioural output, i.e., the choice. When the agent needs to select the best item in a set, it is best to search for evidence that it is more likely to affect the top of the ranking (e.g., is the best item still the best one?); information that changes the middle or the bottom of the ranking is instead less valuable (e.g., is the item ranked as seven is now ranked as six?) because it would not affect the behavioural output. When choosing which item to discard, instead, the optimal strategy involves acquiring information most informative of the *bottom* of the ranking and not the top. We propose that even in the context of binary choice studied here, humans might still deploy this normative strategy, that while it does not provide a normative advantage, it is not suboptimal. Further work in which the size of the set is increased would be required to test this idea more stringently.

The most far reaching conclusion of our work is that context and behavioural demand have a powerful effect on how information is accumulated and processed. Notably, our data show that this is a general effect that spans both more complex value-based choice and simpler perceptual choice. Our conclusion is that, given the limited computational resources of the brain, humans have developed a mechanism that prioritises the processing or recollection of the information that is most relevant for the behavioural response that is required. This has profound implications when we think about the widespread effect of contextual information on decision making that has been at the core of the research in psychology, behavioural economics and more recently neuroeconomics [36, 37, 38, 24, 39]. Most of these contextual or framing effects have been labelled as “biases” because, once one strips away the context, the actual available options should remain identical. However, this perspective may not be putting enough emphasis on the fact that the decision maker has to construct low dimensional (and therefore imperfect) representations of the decision problem. As we have shown here, from the very beginning of the deliberation process, the context — even when it is simple (*like*/*dislike*, *most*/*fewest*) or irrelevant from the experimenter perspective — affects which information is processed, recalled, or attended to, with effects that spread into post-decision processing such as confidence estimation. This, as a consequence, will produce profoundly dissimilar representations according to the behavioural goal set by the context. With this shift of perspective, it may well be that case that many of the so-called “biases” will be shown in a new light, given that participants are dealing with very different choices once the behavioural goal changes. This viewpoint might provide a more encouraging picture of the human mind, by suggesting that evolution has equipped us well to deal with ever-changing environments in the face of limited computational resources.

## 4. Methods

### 4.1 Procedure

#### Value Experiment

At the beginning of this experiment, participants were asked to report on a scale from *£*0-3 the maximum they would be willing to pay for each of 60 snack food items. They were informed that this bid will give them the opportunity to purchase a snack at the end of the experiment, using the Becker-DeGroot-Marschak [40] mechanism, which gives them incentives to report their true valuation. Participants were asked to fast for four hours previous to the experiment, expecting they would be hungry and willing to spend money to buy a snack.

After the bid process, participants completed the choice task: in each trial they were asked to choose between two snack items, displayed on-screen in equidistant boxes to the left and right of the centre of the screen (Figure 1A). After each binary choice, participants also rated their subjective level of confidence in their choice. Pairs were selected using the value ratings given in the bidding task: using a median split, each item was categorized as high- or low-value for the agent; these were then combined to produce 15 high-value, 15 low-value, and 30 mixed pairs, for a total of 60 pairs tailored to the participant’s preferences. Each pair was presented twice, inverting the position to have a counterbalanced item presentation.

The key aspect of our experimental setting is that all participants executed the choice process under two framing conditions: 1) a *like* frame, in which participants were asked to select the item that they liked the most, i.e., the snack that they would prefer to eat at the end of the experiment; and 2) a *dislike* frame in which participants were asked to select the item that they liked the least, knowing that this is tantamount to choosing the other item for consumption at the end of the experiment. See Figure 1A for a diagram of the task.

After 4 practice trials, participants performed a total of 6 blocks of 40 trials (240 trials in total). *Like* and *dislike* frames were presented in alternate blocks and the order was counterbalanced across participants (120 trials per frame). An icon in the top-left corner of the screen (“thumbs up” for *like* and “stop sign” for *dislike*) reminded participants of the choice they were required to make; this was also announced by the investigator at the beginning of every block. The last pair in a block would not be first in the subsequent block.

Participants’ eye movements were recorded throughout the choice task and the presentation of food items was gaze-contingent: participants could only see one item at a time depending on which box they looked at; following Folke and colleagues [7], this was done to reduce the risk that participant, while gazing one item, would still look at the other item in their visual periphery.

Once all tasks were completed, one trial was randomly selected from the choice task. The BDM bid value of the preferred item (the chosen one in the *like* frame and the unchosen one in the *dislike* frame) was compared with a randomly generated number between *£*0-3. If the bid was higher than the BDM generated value, an amount equivalent to the BDM value was subtracted from their *£*20 payment and the participant received the food item. If the bid was lower than the generated value, participants were paid *£*20 for their time and did not receive any snack. In either case, participants were required to stay in the testing room for an extra hour and were unable to eat any food during this time other than food bought in the auction. Participants were made aware of the whole procedure before the experiment began.

#### Perceptual Experiment

Perceptual Experiment had a design similar to the one implemented in Value Experiment, except that alternatives were visual stimuli instead of food items. In this task, participants had to choose between two circles filled with dots (for a schematic diagram see Figure 1), again in two frames. In the *most* frame, they had to pick the one with more dots; and the one with fewer dots in the *fewest* frame. The total number of dots presented in the circles could have three numerosity levels (= 50, 80 and 110 dots). For each pair in those 3 levels, the dot difference between the circles varied in 10 percentage levels (ranging from 2% to 20% with 2% steps). To increase the difficulty of the task, in addition to the target dots (blue-green coloured), distractor dots (orange coloured) were also shown. The number of distractor dots was 80% of that of target dots (40, 64, 88 for the 3 numerosity levels, respectively). Pairs were presented twice and counterbalanced for item presentation. After 40 practice trials (20 initial trials with feedback, last 20 without), participants completed by 3 blocks of 40 trials in the *most* frame and the same number in the *fewest* frame; they faced blocks with alternating frames, with a presentation order counterbalanced across participants. On the top left side of the screen a message indicating *Most* or *Fewest* reminded participants of the current frame. Participants reported their confidence level in making the correct choice at the end of each trial. As in the previous experiment, the presentation of each circle was gaze contingent. Eye tracking information was recorded for each trial. Participants received *£*7.5 for one hour in this study.

Both tasks were programmed using Experiment Builder version 2.1.140 (SR Research).

### 4.2 Exclusion criteria

#### Value Experiment

We excluded individuals that met any of the following criteria:

1. Participants used less than 25% of the BDM value scale.
2. Participants gave exactly the same BDM value for more than 50% of the items.
3. Participants used less than 25% of the choice confidence scales.
4. Participants gave exactly the same confidence rating for more than 50% of their choices.
5. Participants did not comply with the requirements of the experiment (i.e., participants that consistently choose the *preferred* item in *dislike* frame or their average blink time is over 15% of the duration of the trials).

#### Perceptual Experiment

Since for Perceptual Experiment the assessment of the value scale is irrelevant, we excluded participants according to criteria 3, 4 and 5.

### 4.3 Participants

#### Value Experiment

Forty volunteers gave their informed consent to take part in this research. Of these, thirty-one passed the exclusion criteria and were included in the analysis (16 females, 17 males, aged 20-54, mean age of 28.8). To ensure familiarity with the snack items, all the participants in the study had lived in the UK for one year or more (average 17 years).

#### Perceptual Experiment

Forty volunteers were recruited for the second experiment. Thirty-two participants (22 females, 10 males, aged 19-50, mean age of 26.03) were included in the behavioural and regression analyses. Due to instability in parameter estimation, four additional participants were removed from the GLAM modelling analysis.

All participants signed a consent form and both studies were done following the approval given by the University College London, Division of Psychology and Language Sciences ethics committee.

### 4.4 Eye-tracking

#### Value and Perceptual Experiments

An Eyelink 1000 eye-tracker (SR Research) was used to collect the visual data. Left eye movements were sampled at 500 Hz. Participants rested their heads over a head support in front of the screen. Display resolution was of 1024 × 768 pixels. To standardise the environmental setting and the level of detectability, the lighting was monitored in the room using a dimmer lamp and light intensity was maintained at 4±0.5 lx at the position of the head-mount when the screen was black.

Eye-tracking data were analysed initially using Data Viewer (SR Research), from which reports were extracted containing details of eye movements. We defined two interest-areas (IA) for left and right alternatives: two squares of 350 x 350 pixels in Value Experiment and two circles of 170 pixels of radius for Perceptual Experiment. The data extracted from the eye-tracker were taken between the appearance of the elements on the screen (snack items or circle with dots in experiments 1 and 2, respectively) and the choice button press (confidence report period was not considered for eye data analysis).

The time participants spent fixating on each IA was defined the dwelling time (DT). From it, we derived a difference in dwelling time (ΔDT) for each trial by subtracting DT of the right IA minus the DT of the left IA. Starting and ending IA of each saccade were recorded. This information was used to determine the number of times participants alternated their gaze between IAs, i.e., ‘gaze shifts’. The total number of gaze shifts between IAs was extracted for each trial, producing the gaze shift frequency (GSF) variable.

### 4.5 Data Analysis: Behavioural Data

Behavioural measures during *like/dislike* and *most/fewest* frames were compared using statistical tests available in SciPy. All of the hierarchical analyses were performed using lme4 package [41] for R integrated in an Jupyter notebook using the rpy2 package (https://rpy2.readthedocs.io/en/latest/). For choice models, we predicted the log odds ratio of selecting the item appearing at the right. Fixed-effects confidence interval were calculated by multiplying standard errors by 1.96. Additionally, we predicted confidence using a linear mixed-effects model. Predictors were all z-scored at participant level. Matplotlib/Seaborn packages were used for visualization.

### 4.6 Data Analysis: Attentional Model - GLAM

To get further insight on potential variations in the evidence accumulation process due to the change in frames we used the Gaze-weighted Linear Accumulator Model (GLAM) developed by Thomas et al. [11]. GLAM is part of the family of linear stochastic race models in which different alternatives (i, i.e., left or right) accumulate evidence (E_i_) until a decision threshold is reached by one of them, determining the chosen alternative. The accumulator for an individual option is described by the following expression:

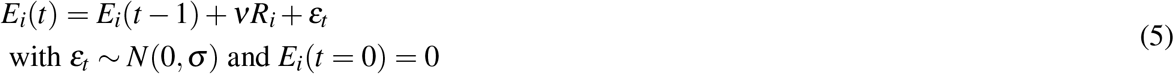

With a drift term (*ν*) controlling the speed of relative evidence (R_i_) integration and i.i.d. noise terms with normal distribution (zero-centered and standard deviation *σ*). R_i_ is a term that expresses the amount of evidence that is accumulated for item i at each time point t. This is calculated as follows. We denote by g_i_, the relative gaze term, calculated as the proportion of time that participants observed item i:

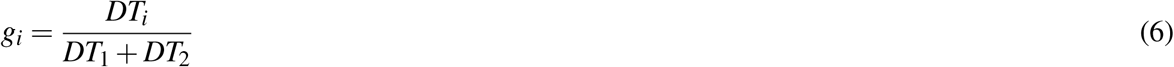

with DT as the dwelling time for item i during an individual trial. Let r_i_ denote the value for item i reported during the initial stage of the experiment. We can then define the average absolute evidence for each item (A_i_) during a trial:

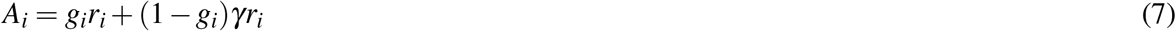

This formulation considers a multiplicative effect of the attentional component over the item value, capturing different rates of integration when the participant is observing item i or not (unbiased and biased states, respectively). The parameter *γ* is the gaze bias parameter: it controls the weight that the attentional component has in determining absolute evidence. Thomas and colleagues [11] interpret *γ* as follows: when *γ* = 1, bias and unbiased states have no difference (i.e., the same r_i_ is added to the average absolute evidence regardless the item is attended or not); when *γ* < 1, the absolute evidence is discounted for the biased condition; when *γ* < 0, there is a leak of evidence when the item is not fixated. Following Thomas et al. [11], in our analysis we allowed *γ* to take negative values, but our results do not change if *γ* is restricted to [0, 1] (Figure S20). Finally, the relative evidence of item i, R_i_*, is given by:

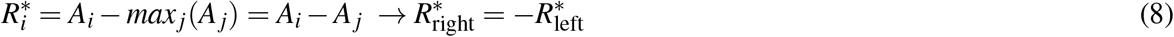

Since our experiment considers a binary choice the original formulation of the model [11], proposed for more than 2 alternatives, R_i_* is reduced to subtract the average absolute evidence of the other item. Therefore, for the binary case, the R_i_* for one item will be additive inverse of the other, e.g., if the left item has the lower value, we would have R_left_* <0 and R_right_* >0. Additionally, in their proposal for GLAM, Thomas and colleagues [11] noted that R_i_* range will depend on the values that the participant reported, e.g., evidence accumulation may appear smaller if participant valued all the items similarly, since R_i_* may be lower in magnitude. This may not represent the actual evidence accumulation process since participants may be sensitive to marginal differences in relative evidence. To account for both of these issues a logistic transformation is applied over R_i_* using a scaling parameter *τ*:

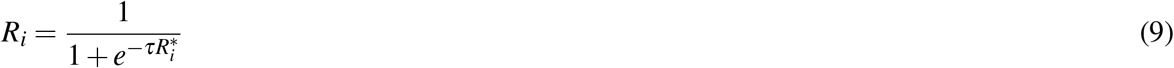

In this case R_i_ will be always positive and the magnitude of the difference between R_right_ and R_left_ will be controlled by *τ*, e.g., higher *τ* will imply a bigger difference in relative evidence (and hence accumulation rate) between left and right item. In the case that *τ* = 0 the participant will not present any sensitivity to differences in relative evidence.

Given that R_i_ represents an average of the relative evidence across the entire trial, the drift rate in E_i_ can be assumed to be constant, which enables the use of an analytical solution for the passage of time density. Unlike aDDM [1], GLAM does not deal with the dynamics of attentional allocation process in choice. Details of these expressions are available at Thomas et al. [11]. In summary, we have 4 free parameters in the GLAM: *ν* (drift term), *γ* (gaze bias), *τ* (evidence scaling) and *σ* (normally distributed noise standard deviation).

The model fit with GLAM was implemented at a participant level in a Bayesian framework using PyMC3 [42]. Uniform priors were used for all the parameters:

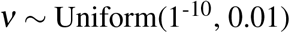

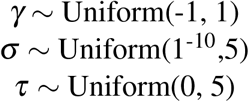

#### Value Experiment

We fitted the model for each individual participant and for *like* and *dislike* frames, separately. To model participant’s behaviour in the *like* frame we used as input for GLAM the RTs and choices, plus BDM bid values and relative gaze for left and right alternatives for each trial. The original GLAM formulation (as presented above) assumes that evidence is accumulated in line with the preference value of a particular item (i.e., “how much I like this item”). When information about visual attention is included in the model, the multiplicative model in GLAM assumes that attention will boost the evidence accumulation already defined by value. Our proposal is that evidence accumulation is a flexible process in which attention is attracted to items based on the match between their value and task-goal (accept or reject) and not based on value alone, as most of the previous studies have assumed. Since in the *dislike* frame the item with the lower value becomes relevant to fulfil the task, we considered the opposite value of the items (r_i,dislike_ = 3 - r_i,like_, e.g., item with value 3, the maximum value, becomes value 0) as an input for GLAM fit. For both conditions, model fit was performed only on even-numbered trials using Markov-Chain-Monte-Carlo sampling, using implementation for No-U-Turn-Sampler (NUTS), 4 chains were sampled, 1000 tuning samples were used, 2000 posterior samples to estimate the model parameters. Model comparison was performed using Watanabe-Akaike Information Criterion (WAIC) scores available in PyMC3, calculated for each individual participant fit.

Pointing to check if the model replicates the behavioural effects observed in the data [43], simulations for choice and response time (RT) were performed using participant’s odd trials, each one repeated 50 times. For each trial, value and relative gaze for left and right items were used together with the individual estimated parameters. Random choice and RT (within a range of the minimum and maximum RT observed for each particular participant) were set for 5% of the simulations, replicating the contaminating process included in the model as described by Thomas et al. [11].

Additionally, we simulated the accumulation process in each trial to obtain a measure of balance of evidence [23, 21] for each trial. The purpose of this analysis was to replicate the effect of ΣValue over confidence (check *Results* for details) and check if it arises from the accumulation process and its interaction with attention. Balance of evidence in accumulator models has been used previously as an approximation to the generation of confidence in perceptual and value-based decision experiments [21, 44, 18]. Consequently, using the value of the items and gaze ratio from odd-numbered trials, we simulated two accumulators (equation 5), one for each alternative. Our simulations used the GLAM parameters obtained from participant’s fit. Once the boundary was reached by one of the stochastic accumulators (fixed boundary = 1), we extracted the simulated RT and choice. The absolute difference between the accumulators when the boundary was reached (Δe = |E_right_(t_final_) - E_left_(t_final_)|) delivered the balance of evidence for that trial. In total 37200 trials were simulated (10 repetitions for each one of the trials done by the participants). A linear regression model to predict simulated Δe using |ΔValue|, simulated RT and ΣValue as predictors was calculated with the pooled participants’ data. This model was chosen since it was the most parsimonious model obtained to predict participant’s confidence in the Value Experiment (Figure S13). The best model includes GSF as predictor in the regression, but since GLAM does not consider the gaze dynamics we removed it from the model. Δe simulations using a GLAM without gaze influence (i.e., equal gaze time for each alternative) were also generated, to check if gaze difference was required to reproduce ΣValue effect over confidence. The parameters fitted for individual participants were also used in the no-gaze difference simulation. The same linear regression model (Δe ~ | ΔValue| + simulated RT + ΣValue) was used with the data simulated with no-gaze difference.

#### Perceptual Experiment

In the Perceptual Experiment, we repeated the same GLAM analysis done in Value Experiment. Due to instabilities in the parameters’ fit for some participants, we excluded 4 extra participants. Twenty-eight participants were included in this analysis. Additionally, the GLAM fit in this case was done removing outlier trials, i.e., trials with RT higher than 3 standard deviations (within participant) or higher than 20 seconds. Overall less than 2% of the trials were removed. For *most* frame, relative gaze and perceptual evidence (number of dots) for each alternative were used to fit choice and RT. In a similar way to the consideration taken in the *dislike* case, we reassigned the perceptual evidence in the *fewest* frame (r_i,fewest_ = 133 - r_i,most_+ 40, considering that 133 is the higher number of dots presented and 40 dots the minimum) in a way that the options with higher perceptual evidence in the *most* frame have the lower evidence in the *fewest* frame. The same MCMC parameters used to fit the model for each participant in the Value Experiment were used in this case (again, only even-numbered trials were used to fit the model). Behavioural out-of-sample simulations (using the odd-numbered trials) and balance of evidence simulations (33600 trials simulated in the Perceptual Experiment) were considered in this analysis. We tested the effect of ΣDots over confidence with a similar linear regression model than the one used in the Value Experiment. Pooled participants’ data for ΔDots, simulated RT and ΣDots was used to predict Δe. Δe simulations using a GLAM without gaze asymmetry were also calculated in this case. All the figures and analysis were done in python using GLAM toolbox and custom scripts.

## Acknowledgments

This study was funded by a Sir Henry Dale Fellowship (102612 /A/13/Z) awarded to Benedetto De Martino by the Wellcome Trust. Pradyumna Sepulveda was funded by the Chilean National Agency for Research and Development (ANID)/Scholarship Program/DOCTORADO BECAS CHILE/2017 - 72180193. We would like to thank Antonio Rangel for his valuable comments on an earlier version of the manuscript and Mariana Zurita for the help in the proofreading of the manuscript.

## Data Availability

All the data and the codes used for this study will be made available upon acceptance of the manuscript.

