## Supplemental Information for "Visual attention modulates the integration of goal-relevant evidence and not value"

Pradyumna Sepulveda[1]*, Marius Usher[2], Ned Davies[1], Amy Benson[1], Pietro Ortoleva[3] and Benedetto De Martino[1,4*].

1. Institute of Cognitive Neuroscience, University College London, London, United Kingdom.
2. School of Psychological Sciences and Sagol School of Neuroscience, Tel Aviv University, Tel Aviv, Israel.
3. Department of Economics and Woodrow Wilson School, Princeton University.
4. Wellcome Centre for Human Neuroimaging, University College London, London, United Kingdom.

### SUPPLEMENTAL INFORMATION

**INDEX**


SI 1 : Task Framing Differences 3

SI 2 : Choice Regression Models 9

SI 3 : Fixation Analysis 14

SI 4 : Confidence Regression Models 19

SI 5: GLAM - Model Comparison and Out-of-Sample Simulations 21

SI 6: GLAM - Parameter Comparison 26

SI 7: Attentional Drift Diffusion Model 29

SI 8: GLAM - Balance of Evidence Simulations 34

SI 9: Normative Model - Proof of Propositions 1 and 2 37

Supplemental References 40

### SI 1: Task Framing Differences

*Value Experiment.* We examined how the frame manipulation impacted overall performance (Figure S1A). We defined "accuracy" as the proportion of trials in which participant's reported values (BDM bid) correctly predicted their binary decision, i.e., they select the item with highest value in the *like* frame and the one with lowest value in the *dislike* frame. Overall accuracy was not significantly different in both frames ($Mean_{Like}$=0.77; $Mean_{Dislike}$=0.75, t=1.71; p=0.1). We also found that participants had slightly slower reaction times (RTs) in the *dislike* frame ($Mean_{Like}$=2858.2 ms, $Mean_{Dislike}$=3152.7 ms; t=-2.52; p<0.05). Participants reported lower confidence in the *dislike* frame ($Mean_{\Delta Confidence}$=0.19; t=4.49; p<0.001) and shifted their gaze (gaze shift frequency, GSF) between items more during *dislike* trials ($Mean_{\Delta|GSF|}$=-0.110; t=-2.99; p<0.01). These results overall suggest that the subjects may have found the *dislike* condition slightly less intuitive. Although this did not affect their performance, it slightly reduced their confidence and increased RT and GSF.

As observed in previous studies [1,2], we found that choice accuracy was modulated by confidence: decisions in which participants reported high-confidence were more accurately predicted by the value estimate collected before the experiment – the slope of the logistic curve is steeper in the case of high confidence (Figure 1B, *Results* section). In this study, this effect is replicated in both *like* (low confidence: β=0.769; high confidence: β=1.633) and *dislike* (low confidence: β=-0.642; high confidence: β=-1.363) frames. Note that the inversion of the sign of the slopes in *like* vs *dislike* frames indicate that participants were performing the task correctly ($\Delta\beta_{Like-Dislike}$: t=8.14, p<0.001), selecting the item with lower value during the *dislike* frame (Figure 1C, *Results* section). Choice accuracy (steepness of the slopes) was not significantly different between *like* and *dislike* frames ($\Delta|\beta_{Like-Dislike}|$: t=1.58, p=0.124).

*Perceptual Experiment*. We repeated the same analysis for the behavioural performance in *most* and *fewest* frames (Figure S1B). In contrast to the Value Experiment, we observed a slight reduction in accuracy in participant responses for the *fewest* frame ($Mean_{Most}$=0.77, $Mean_{Few}$=0.74, t=2.46; p<0.05); unlike the Value Experiment, however, we did not find differences in RTs ($Mean_{Most}$=4029.57 ms, $Mean_{Few}$=3975.59 ms; t=0.32; p=0.75). During the *fewest* frame participants reported lower confidence ($Mean_{\Delta Confidence}$=0.24; t=5.62; p<0.001) and shifted their gaze more between alternatives ($Mean_{\Delta|GSF|}$=-0.17; t=-4.15; p<0.001), as observed in the Value Experiment.

Participants also reported higher confidence in trials that better discriminated the number of dots (Figure 1E, *Results* section). This effect was replicated in both *most* (low confidence: β=1.142; high confidence: β=2.164) and *fewest* (low confidence: β=-1.118; high confidence: β=-2.010) frames. The inversion of the sign of the slopes in *most* vs *fewest* frames also shows that participants were performing correctly ($\Delta\beta_{\text{Most-Few}}$: $t = 22.22$, $p<0.001$); the magnitude of the slopes was not significantly different between the two frames ($\Delta|\beta_{\text{Most-Few}}|$: t=0.79, p=0.434; Figure 1F, *Results* section). This pattern of results mirrors the pattern seen in the Value Experiment.

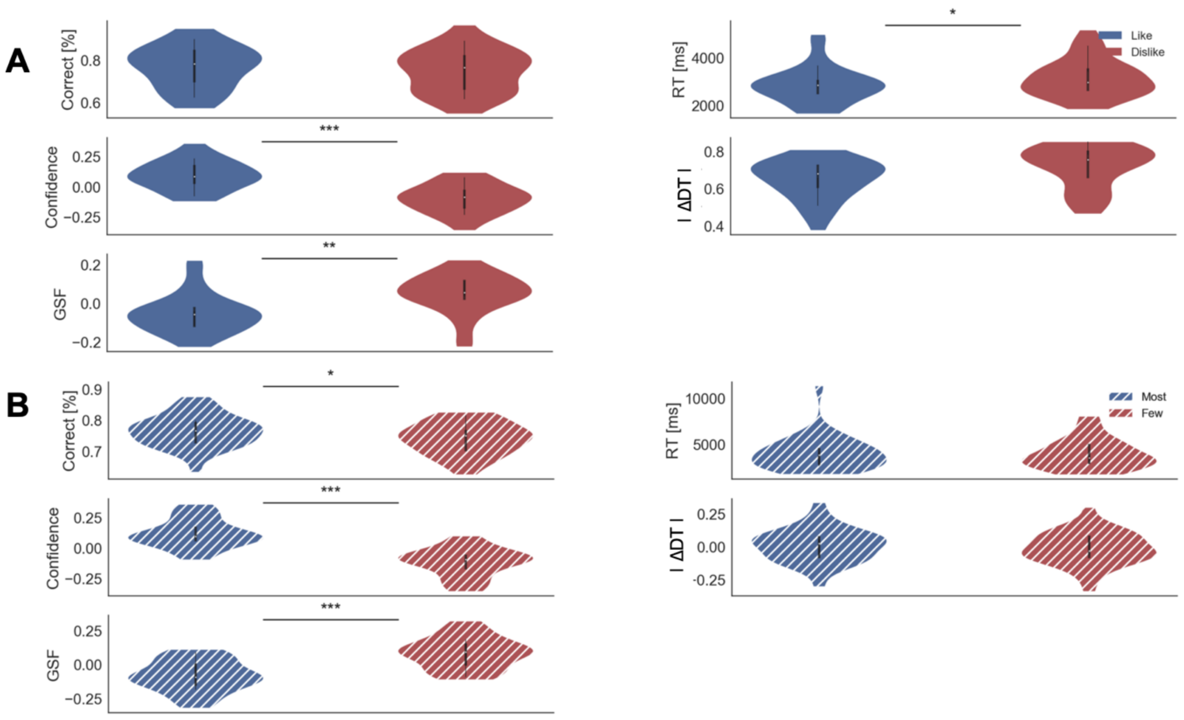


*Figure S1. Behavioural results for Value (A) and Perceptual (B) Experiments. Confidence, DDT and GSF values have been z-scored per participant. In the violin plot, red and blue areas indicate the distribution of the parameters across participants. Black bars present the 25, 50 and 75 percentiles of the data. Solid colour indicates the Value Experiment and striped colours indicate the Perceptual Experiment. RT: reaction time; ∆DT: Difference in Dwell Time; GSF: Gaze Shift Frequency.*

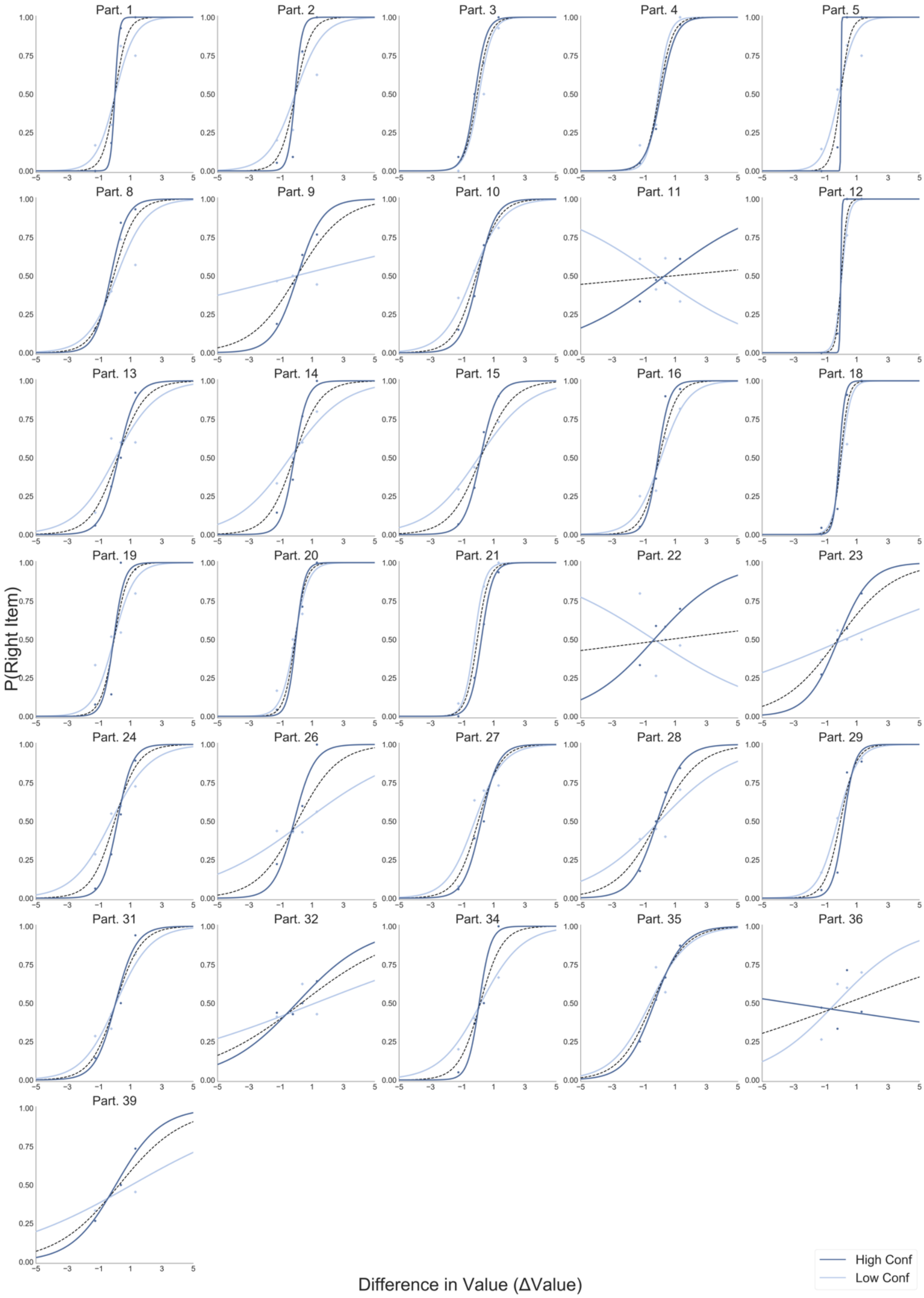


*Figure S2. Logistic regression predicting choice from the difference in value between the two items (∆Value). All participants in the Value Experiment, like frame, are presented. Light blue lines depict the logistic fit calculated using only low confidence trials. Dark blue lines show the logistic fit only for high confidence trials. Segmented black line considers the logistic regression calculated using all the trials.*

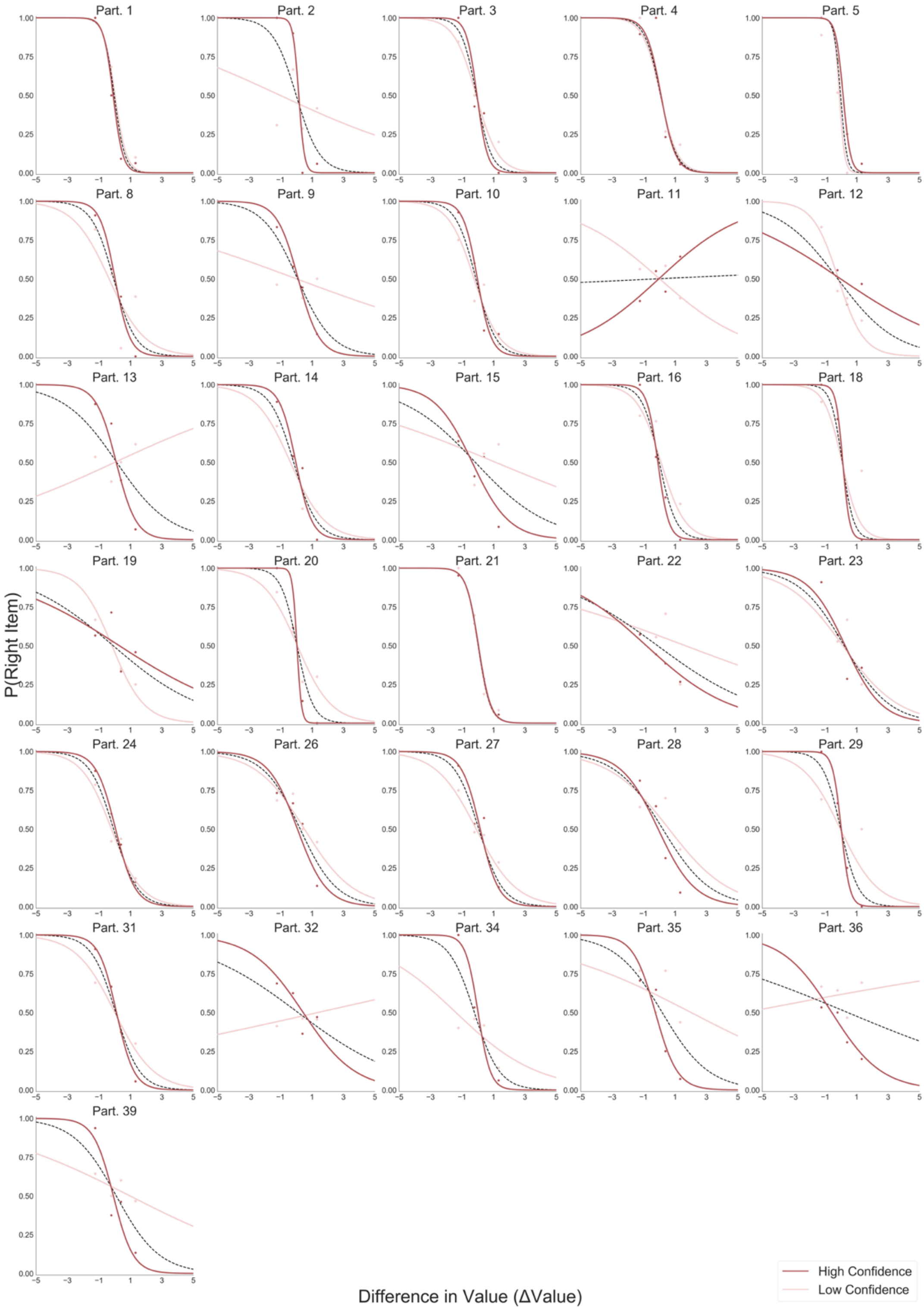


*Figure S3. Logistic regression predicting choice from the difference in value between the two items (ΔValue). All participants in the Value Experiment, dislike frame, are presented. Light red lines depict the logistic fit calculated using low confidence trials. Dark red lines show the logistic fit using high confidence trials. Segmented black line considers the logistic regression calculated with all the trials.*

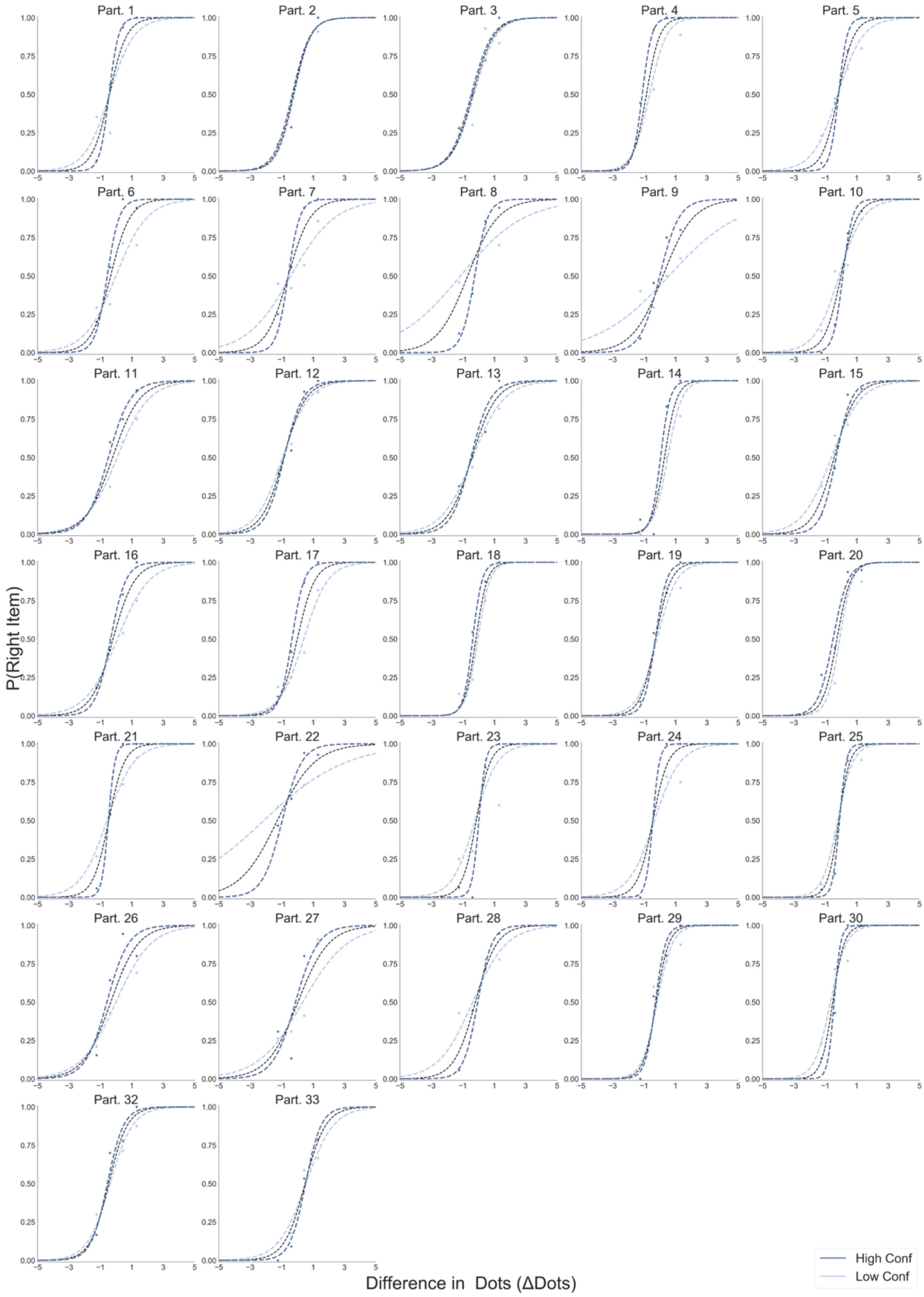


*Figure S4. Logistic regression predicting choice from the difference in number of dots between the two circles (ΔDots). All participants in the Perceptual Experiment, most frame, are presented. Light blue lines depict the logistic fit calculated using only low confidence trials. Dark blue lines show the logistic fit only for high confidence trials. Segmented black line considers the logistic regression calculated with all the trials.*

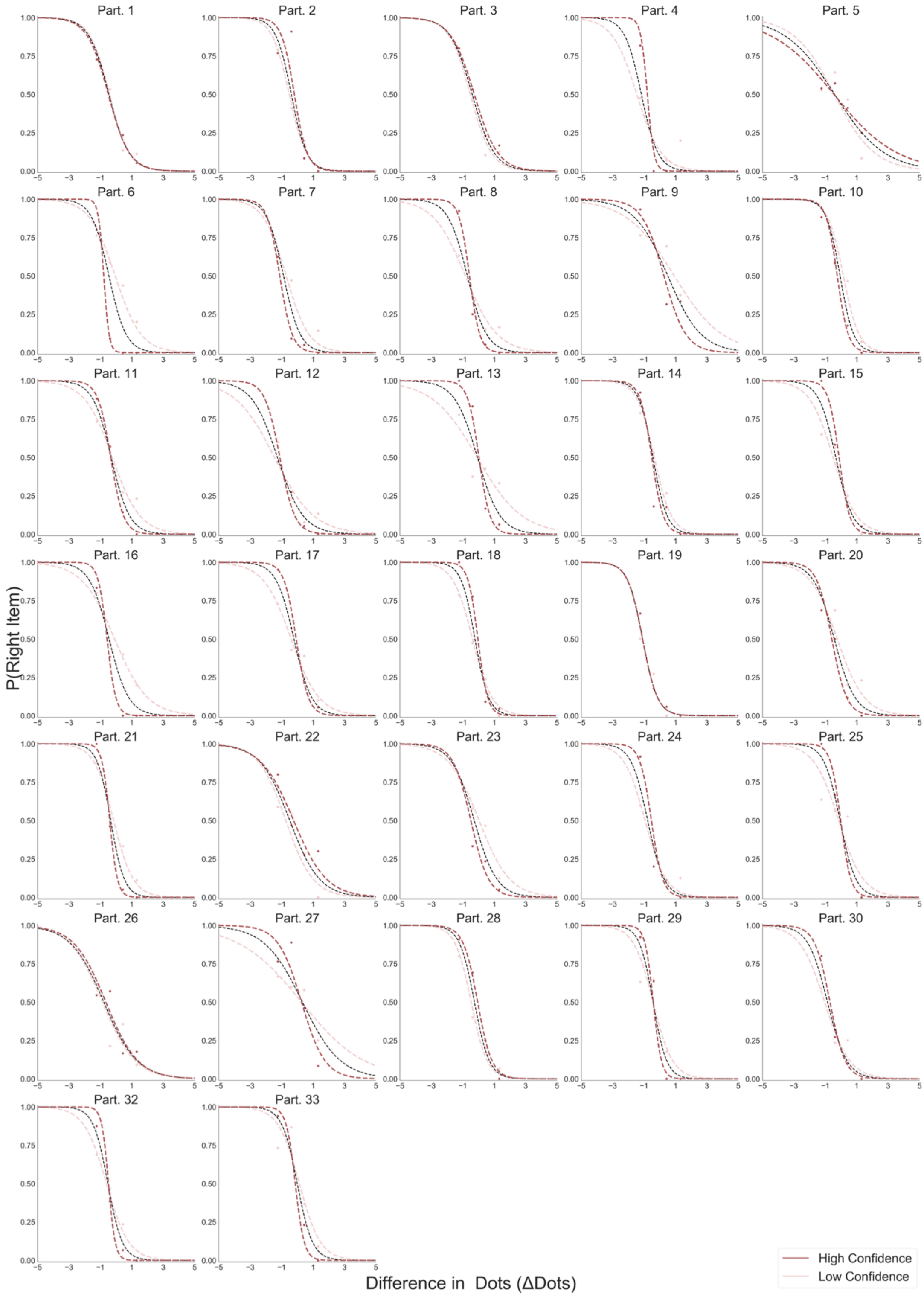


*Figure S5. Logistic regression predicting choice from the difference in number of dots between the two circles (ΔDots). All participants in the Perceptual Experiment, fewest frame, are presented. Light red lines depict the logistic fit calculated using only low confidence trials. Dark red lines show the logistic fit only for high confidence trials. Segmented black line considers the logistic regression calculated with all the trials.*

### SI 2: Choice Regression Models

*Table S1. Hierarchical logistic models for choice*

| Models | Formulas |
|---|---|
| Model 1 | Choice ~ ∆Value |
| Model 2 | Choice ~ ∆Value + Confidence |
| Model 3 | Choice ~ ∆Value + Confidence + ΣValue |
| Model 4 | Choice ~ ∆Value + Confidence + ΣValue + ∆DT |
| Model 5 | Choice ~ ∆Value + Confidence + ΣValue + ∆DT + ∆Value * Confidence |
| Model 6 | Choice ~ ∆Value + Confidence + ΣValue + ∆DT + ∆Value * Confidence + ∆Value * ΣValue |
| Model 7 | Choice ~ ∆Value + Confidence + ΣValue + ∆DT + ∆Value * Confidence + ∆Value * ΣValue + Confidence * ∆DT |
| Model 8 | Choice ~ ∆Value + Confidence + ΣValue + ∆DT + GSF + ∆Value * Confidence + ∆Value * ΣValue + Confidence * ∆DT + ∆Value * GSF |

In Value Experiment: ∆Value: difference in value; ΣValue: summed value; ∆DT: difference in dwell time; GSF: gaze shift frequency. In Perceptual Experiment similar models were compared but replacing ∆Value for ∆Dots and ΣValue for ΣDots.

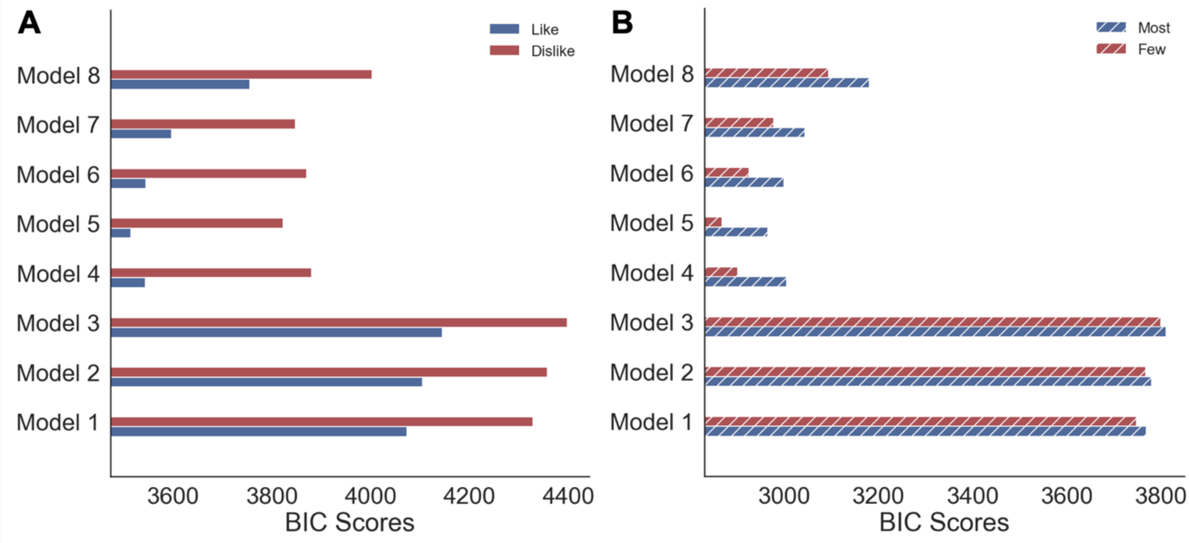


*Figure S6. Model comparison of hierarchical logistic regressions for choice. (A) Value and (B) Perceptual Experiments. Solid colour indicates the Value Experiment and striped colours indicate the Perceptual experiment.*

*Value Experiment*. Using a logistic hierarchical regression model, we investigated which factors modulated choice-proportion, defined here as the probability of choosing the item on the right side of the screen. We report here the results of the most parsimonious model (i.e., the model with a lowest BIC; Figure S6) fitted to the *like* and *dislike* frames independently (Figure 2B, *Results* section). In Table S1 we present the parameters for each factor included in the model. In the *like* frame, the difference in the value of the right item minus left item (ΔValue) had a positive influence on choice-proportion, i.e., participants selected the items that had higher value. This is reversed in the *dislike* frame: ΔValue is now a *negative* predictor of choice, i.e., participants selected the items that had lower value. In both conditions, confidence enhanced the effect of ΔValue, as shown by the interaction between ΔValue and confidence in the *like* and *dislike* frame. These results confirm the findings presented in Figure 1B (*Results* section) while controlling for other relevant variables. Unsurprisingly, confidence and summed value (ΣValue, the added value of both alternatives) were found to show no main effect on the choice-proportion. As discussed in the Results section, gaze allocation (difference in dwell time, ΔDT) is directed to the chosen item in both frames, i.e., the parameters are positive for ΔDT in *like* and *dislike* frame.

*Table S2. Statistical results for the hierarchical linear models for choice in Value Experiment. Z-values for the regression coefficients and their statistical significance are presented for both frames. To check significant differences of the regression coefficients between like and dislike frames repeated samples t-tests between the participants' regression coefficients were calculated.*

| | Choice Value Experiment | | | | | |
|---|---|---|---|---|---|---|
| | Like | | Dislike | | Like - Dislike | |
| | z | p | z | p | t | p |
| ∆Value | 7.917 | <0.001 | -8.652 | <0.001 | 10.74 | <0.001 |
| ∆DT | 6.448 | <0.001 | 6.75 | <0.001 | 2.31 | <0.05 |
| ∆Value x Conf | 5.446 | <0.001 | -4.681 | <0.001 | 9.55 | <0.001 |

* Confidence and ΣValue did not have a significant effect over choice in the regression.

*Perceptual Experiment*. As in the Value Experiment, we used a logistic hierarchical regression to determine the relevant factors modulating perceptual choice (choosing the circle with dots on the right side of the screen) (Figure 2D, *Results* section). We found that the most parsimonious model for choice was the same used in the Value Experiment, where *like* and *dislike* were replaced by *most* and *fewest* frames (Figure S6B). In the *most* frame, the difference in the number of dots of the right alternative minus the left one (∆Dots) had a positive influence over choice; that is, participants tended to select the circle with more dots. As expected, this pattern was reversed in the *fewest* frame: ∆Dots was a negative predictor of choice. As in the Value Experiment, confidence modulated the effect of ∆Dots in *most* and *fewest* frames. The sum of dots presented in both circles during a trial (ΣDots) was found not to have a significant effect on either frame, as expected. However, as discussed in the *Results* section, confidence was found to be a negative predictor of choice in *most* and *fewest* frames. This means participants had a bias to report higher confidence when they chose the left circle. In a similar way to the Value Experiment, participants spend more time fixating the chosen alternative in both frames, with ∆DT effect being positive in *most* and *fewest* frames.

*Table S3. Statistical results for the hierarchical logistic models for choice in Perceptual Experiment. Z-values for the regression coefficients and their statistical significance are presented for both frames. Repeated samples t-tests between the participants' regression coefficients in most and fewest frames were calculated.*

| | Choice Perceptual Experiment | | | | | |
|---|---|---|---|---|---|---|
| | Most | | Fewest | | Most - Fewest | |
| | z | p | z | p | t | p |
| ∆Dots | 14.905 | <0.001 | -14.394 | <0.001 | 30.32 | <0.001 |
| Confidence | -2.823 | <0.01 | -6.705 | <0.001 | 6.67 | <0.001 |
| ∆DT | 10.249 | <0.001 | 10.449 | <0.001 | -2.17 | <0.05 |
| ∆Dots x Conf | 8.677 | <0.001 | -6.23 | <0.001 | 23.69 | <0.001 |
| * ΣDots did not have a significant effect over choice in the regression. | | | | | | |

In a study by Kovach and colleagues [3] a design similar to our value-based experiment was implemented. Participants were required to indicate the item to keep and the one to discard. They found, similarly to our findings in the value-based study, that the overall pattern of attention was mostly allocated according the task goal. However, in the first few hundred milliseconds, these authors found that attention was directed more prominently to the most valuable item in both conditions. We did not replicate this last finding in our experiment, but one possible reason for this discrepancy is that the experiment by Kovach and colleagues presented both items on the screen at the beginning of the task -- unlike in our task, in which the item was presented in a gaze-contingent way (to avoid processing in the visual periphery). This setting might have triggered an initial and transitory bottom-up attention grab from the most valuable (and often most salient) item before the accumulation process started.

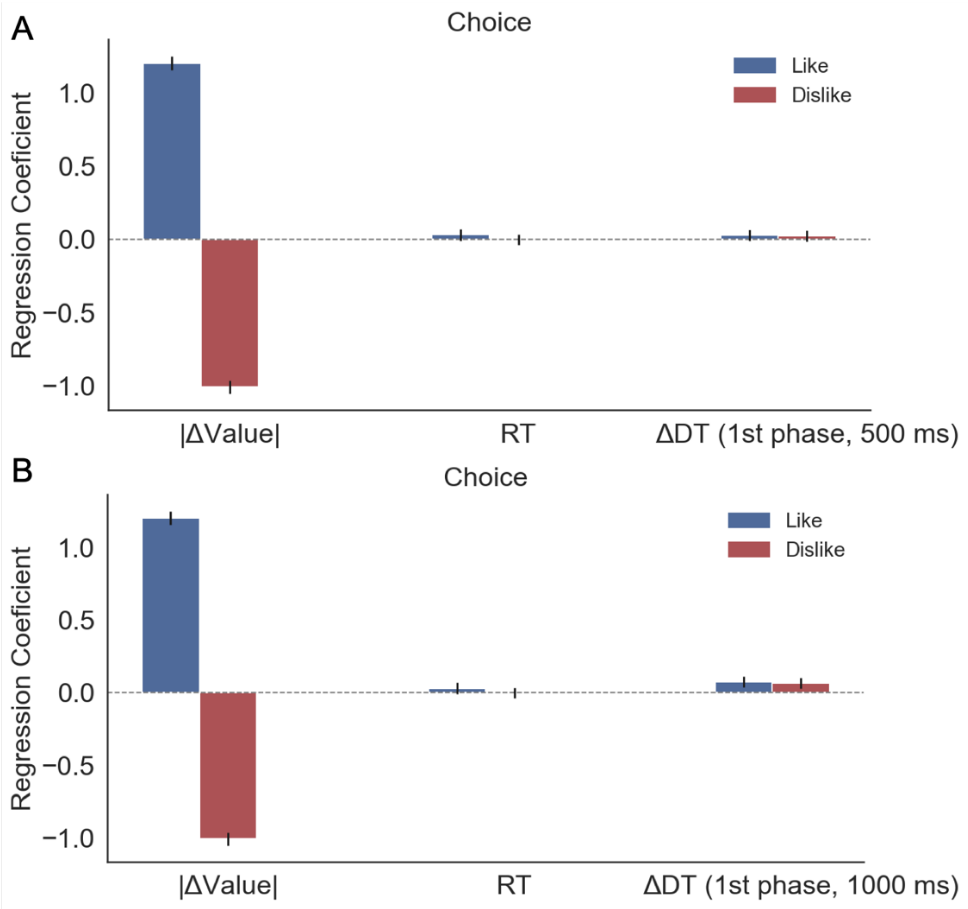


*Figure S7. Kovach and colleagues [3] conducted a study in which participants have to choose food items in 'keep' and 'discard' frames, in a similar way to our Value Experiment. Gaze allocation was found to gravitate towards the chosen item overall, although during the initial moments of the trial (≈ 500 ms), they reported that gaze was directed towards the preferred item. To check if this effect appears in our Value Experiment we ran a regression model to predict choice (i.e., probability of choosing the item presented on the right side of the screen). We restricted the time to estimate ∆DT to the first 500 ms of the trial and used that variable as a predictor of choice in our model (A). We did not find a significant effect of gaze over choice in that period. This difference may be caused by the way the alternatives were presented during the decision time: while in Kovach et al. [3] both alternatives were always displayed on screen during deliberation time, in our experiment the presentation was gaze contingent (i.e., participants needed to explore both items at the beginning of the trial to identify the available items). (B) We recalculated the model considering the initial 1000 ms of the trial and we observe how ∆DT starts to increase its effect over choice. The positive effect of ∆DT over choice is only significant ($z = 1.97$, $p<0.05$) in the like frame; in dislike frame the small effect is only a trend ($z = 1.081$, $p = 0.07$). However, at 1000ms ∆DT is already starting to be allocated to the option coherent with the behavioural responses required by the frame, not to preference.*

### SI 3: Fixation Analysis

In the main text we reported the analysis of last fixation and how its allocation to the (chosen) goal-relevant alternative is modulated by value/number of dots. This result confirmed the findings in Krajbich et al.[4] and expanded them to *dislike* frame and the perceptual realm. To give a more complete view of the fixations properties we additionally performed a similar analysis to Krajbich et al.[4] for first and middle fixations.

It is important to notice that in our Value and Perceptual Experiments, at the beginning of each trial participants do not visualize the options since the presentation is gaze contingent. Therefore, an initial exploration is required to identify the alternatives involved in the decision. In Krajbich et al. [4] both options are visible from the beginning of the trial, however, participants' initial fixation is still randomly allocated.

For the analysis of middle fixations, if blank fixations were recorded between fixations to the same item, then those fixations were assigned to that item (e.g. 'Right', 'Blank', 'Right' was considered as 'Right', 'Right', 'Right'). Trials without middle fixations (i.e. only a first and a last fixation) were removed from the analysis. Trials with no item fixations for more than 40ms at the beginning of the trial were also removed. In the following figures, results from Krajbich et al. [4] are presented together with our findings, as a reference.

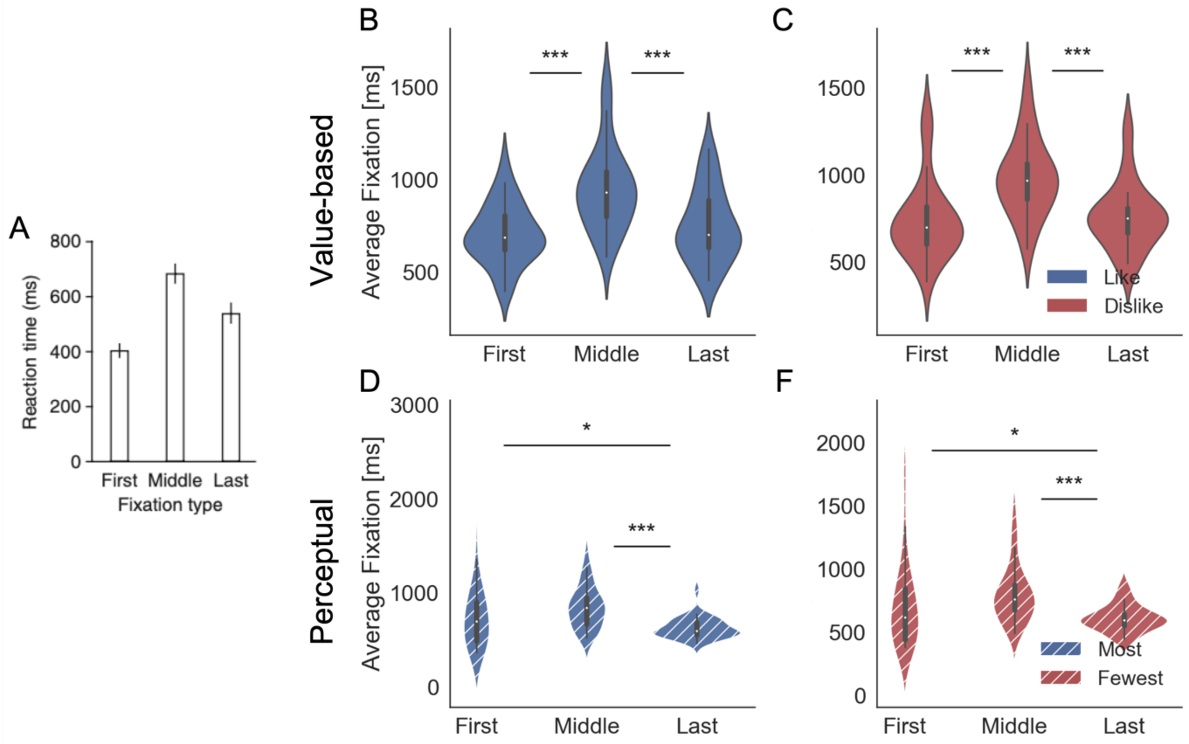


*Figure S8. Fixation duration by type. Middle fixations indicate any fixations that were not the first or last fixations of the trial. (A) In Krajbich et al. [4] middle fixations were found to be longer than first and last fixations on average. For our Value Experiment, in like (B) and dislike (C)*

*frames, and Perceptual Experiment, in most (D) and fewest (F) frames, the same pattern emerges with middle usually longer that first and last fixations. Violin plots depict the distribution of participant's average fixation time. Panel A reproduced from Krajbich et al. [4] .***: p < 0.001, **: p < 0.01, *: p < 0.05.*

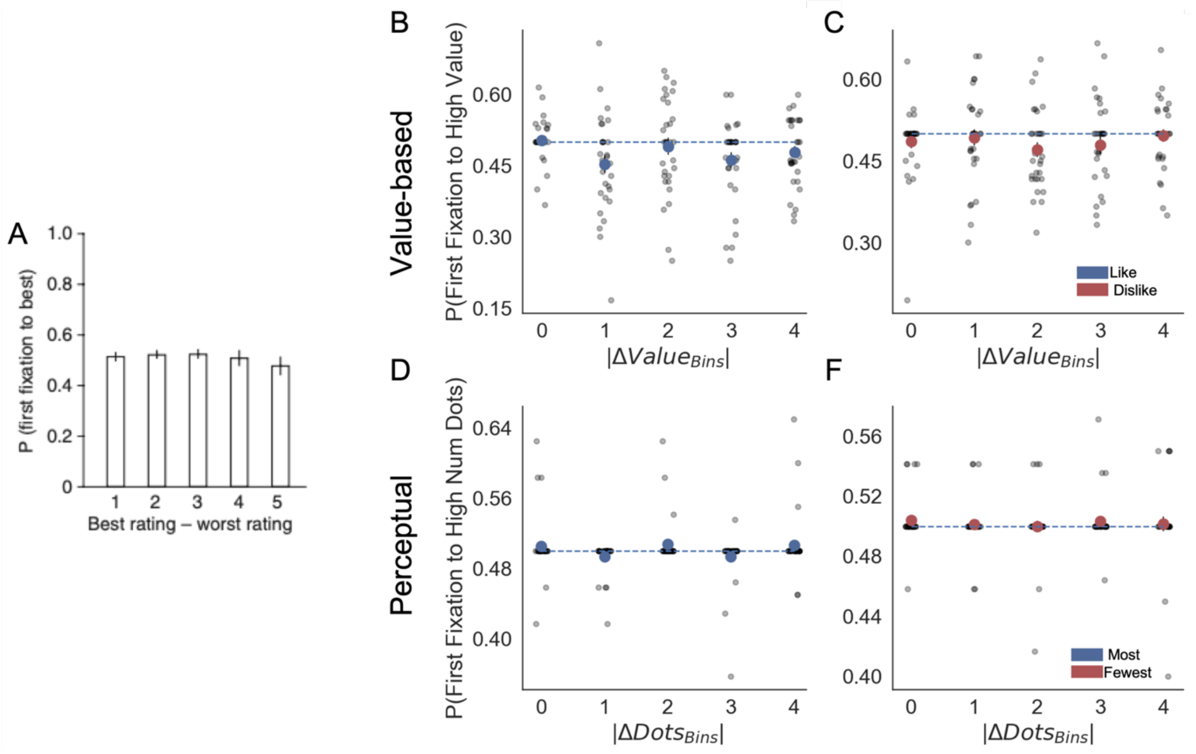


*Figure S9. Fixation properties: probability that the first fixation is to the best item. (**A**) Krajbich et al. [4] reported that the probability is not significantly different from 50%, unaffected by the difference in ratings or difficulty (in our experiments difficulty is equivalent to the absolute difference item value, |ΔValue|, and absolute difference in number of dots, |ΔDots|) . A similar pattern emerges in our Value Experiment, for like (B) and dislike (C) frames, and Perceptual Experiment, for most (D) and fewest (F) frames. Participant responses did not diverge from chance. Importantly, while in Krajbich et al. [4] participants can see both alternatives from the beginning of the trial, our presentation was gaze contingent. Segmented blue line indicates chance level. Light grey dots correspond to individual participants' probability of first fixation to high value/number of dots alternatives for each bin. Red or blue circles indicate the average for that bin considering all the participants. Panel A reproduced from Krajbich et al. [4].*

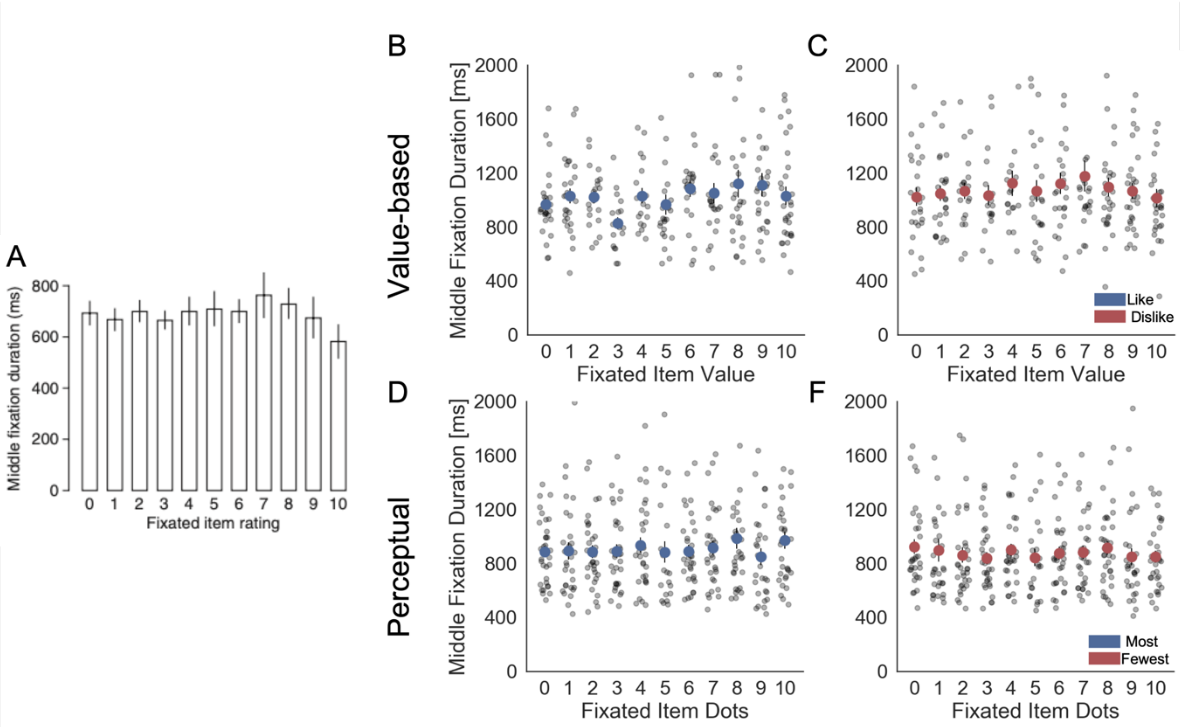


*Figure S10. Fixation properties: middle fixation duration as a function of the rating (value or number of dots) of the fixated item. (A) Krajbich et al.[4] reported that middle fixations durations were independent of the value of the fixated items. In Value Experiment, we found that middle fixation duration was independent of the value of the fixated item in like frame (B), however a slight yet significant effect in dislike (C) frame was found (hierarchical linear regression estimate: $\beta_{Dislike} = 0.025$, $t = 3.465$, $p < 0.001$). In the Perceptual Experiment, for the most (D) frame we found a significant effect of fixated value ($\beta_{Most} = 0.018$, $t = 3.066$, $p < 0.01$), but not for fewest (F) frame. Light grey dots correspond to individual participants' middle fixation durations for each bin. Red or blue circles indicate the average for that bin considering all the participants. For the hierarchical linear regression z-scored data at participant levels was used. Panel A reproduced from Krajbich et al. [4].*

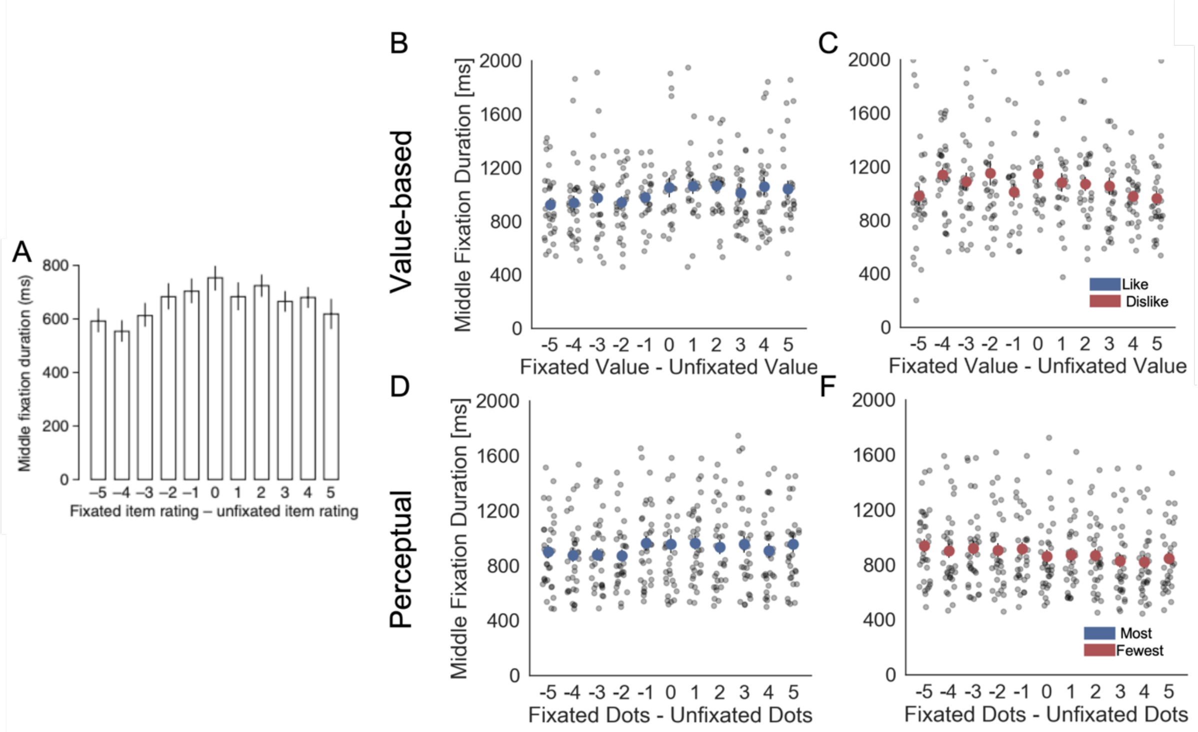


*Figure S11. Fixation properties: middle fixation duration as a function of the difference in liking ratings (value or number of dots) between the fixated and unfixated items. (A) Krajbich et al.[4] reported a slight but significant dependency of middle fixations durations on the difference in value between items. In our Value Experiment, we found that in like and dislike this relationship was significant (hierarchical linear regression estimate: $\beta_{Llike}$ = 0.013, t = 2.031, p<0.05; $\beta_{Dislike}$=-0.027, t=-4.469, p<0.001). Similarly, in the Perceptual Experiment the dependence was found also significant ($\beta_{Most}$ = 0.01, t = 2.551, p <0.05; $\beta_{Few}$=-0.027, t=6.67, p<0.001). Interestingly, a positive sign of the effect in like and most frames would indicates that middle fixations tend to be longer for the option with the higher value or number of dots. On the other hand, a negative sign indicates that middle fixations would be longer for the option with lower value or number of dots in dislike and fewest frames. Light grey dots correspond to individual participants' middle fixation durations for each bin. Full red or blue circles indicate participant's average. Data is binned across participants for visualization. Panel A reproduced from Krajbich et al. [4].*

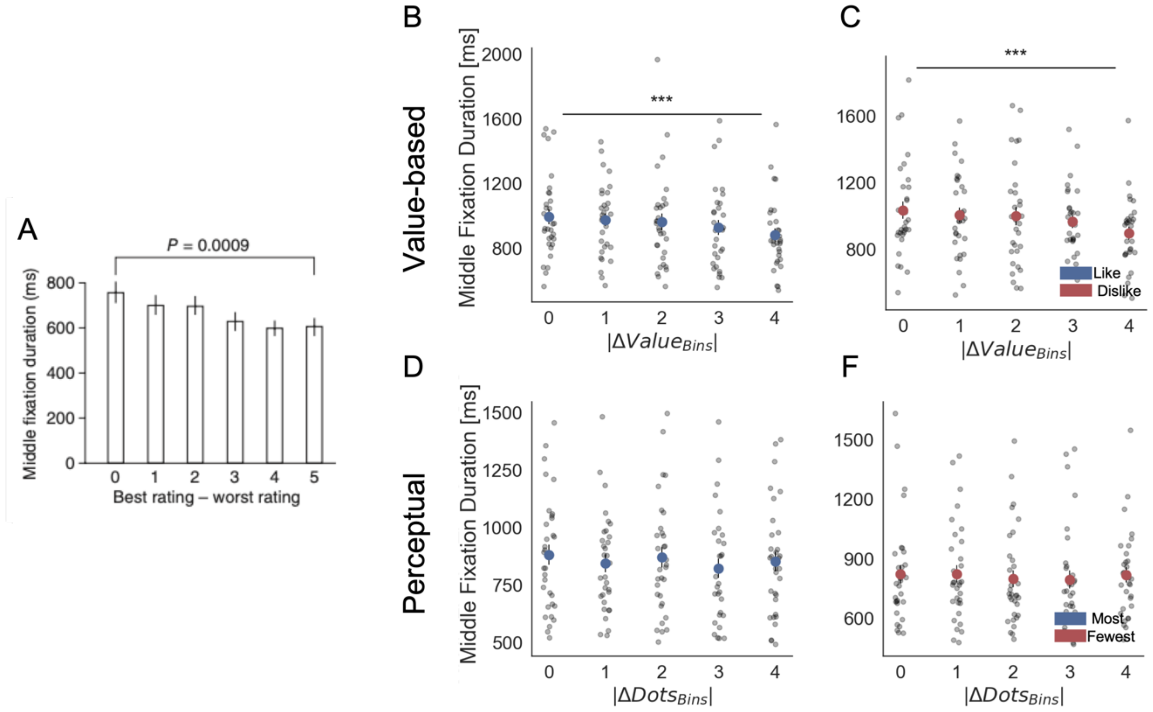


*Figure S12. Fixation properties: middle fixation duration as a function of the difference in ratings between the best- and worst-rated items (difficulty of the trial). In our experiments |ΔValue| and |ΔDots| represent the difficulty of the trials. (A) Krajbich et al.[4] reported a dependency of middle fixations durations on difficulty, with longer fixations in more difficult decisions. In our Value Experiment, in like and dislike frames a similar pattern was found: longer middle fixations for more difficult (lower |ΔValue|) trials (hierarchical linear regression estimate: $\beta_{Llike}$ = -0.053, t = -3.138, p<0.01; $\beta_{Dislike}$=-0.079, t=-5.128, p<0.001). The same relationship was found only in the most frame in the Perceptual Experiment ($\beta_{Most}$ = -0.051, t = -5.128, p <0.01; $\beta_{Few}$=-0.005, t=-0.359, p=0.71). Light grey dots correspond to individual participants' middle fixation durations for each bin. Full red or blue circles indicate participant's average. Data is binned across participants for visualization. Panel A reproduced from Krajbich et al. [4]. Tests presented here are based on a paired two-sided t-test between the first and last bin.***: p < 0.001, **: p < 0.01, *: p < 0.05*

### SI 4: Confidence Regression Models

*Table S4. Hierarchical linear models for confidence*

| **Models** | **Formulas** |
|---|---|
| Model 1 | Confidence ~ \|ΔValue\| |
| Model 2 | Confidence ~ \|ΔValue\| + RT |
| Model 3 | Confidence ~ \|ΔValue\| + RT + GSF |
| Model 4 | Confidence ~ \|ΔValue\| + RT + GSF + ΣValue |
| Model 5 | Confidence ~ \|ΔValue\| + RT + GSF + ΣValue + ΔDT |

In Value Experiment: |ΔValue|: absolute difference in value; RT: reaction time; ΣValue: summed value; ΔDT: difference in dwell time; GSF: gaze shift frequency. In Perceptual Experiment similar models were compared, but replacing ΔValue for ΔDots and ΣValue for ΣDots.

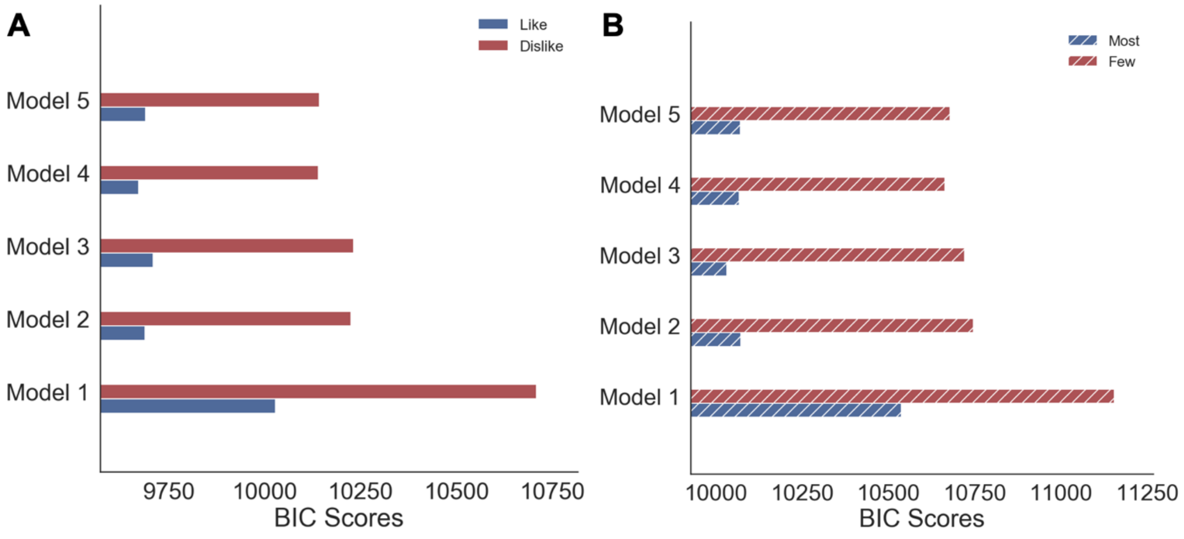


*Figure S13. Model comparison of hierarchical linear regressions for confidence. (A) Value and (B) Perceptual Experiments. Solid colour indicates the value-based experiment and striped colours indicate the perceptual experiment.*

*Table S5. Statistical results for the hierarchical linear models for confidence in Value Experiment. Z-values for the regression coefficients and their statistical significance are presented for the two frames. Repeated samples t-tests between the participants' regression coefficients in like and dislike frames were calculated.*

| | Confidence Value Experiment | | | | | |
|---|---|---|---|---|---|---|
| | Like | | Dislike | | Like - Dislike | |
| | z | p | z | p | t | p |
| \|ΔValue\| | 5.465 | <0.001 | 6.3 | <0.001 | -4.72 | <0.01 |
| RT | -6.373 | <0.001 | -7.739 | <0.001 | ns | |
| GSF | -2.365 | <0.05 | -2.589 | <0.05 | ns | |
| ΣValue | 3.206 | <0.001 | -4.492 | <0.001 | 9.91 | <0.001 |

*Table S6. Statistical results for the hierarchical linear models for confidence in Perceptual Experiment. Z-values for the regression coefficients and their statistical significance are presented for the two frames. Repeated samples t-tests between the participants' regression coefficients in most and fewest frames were calculated.*

| | Confidence Perceptual Experiment | | | | | |
|---|---|---|---|---|---|---|
| | Most | | Fewest | | Most - Fewest | |
| | z | p | z | p | t | p |
| \|ΔValue\| | 3.546 | <0.001 | 7.571 | <0.001 | -4.554 | <0.001 |
| RT | -7.599 | <0.001 | -5.51 | <0.001 | ns | |
| GSF | -4.354 | <0.001 | -5.204 | <0.001 | ns | |
| ΣDots | *2.061* | <0.05 | -7.135 | <0.001 | 14.621 | <0.001 |

### SI 5: GLAM – Model Comparison and Out-of-Sample Simulations

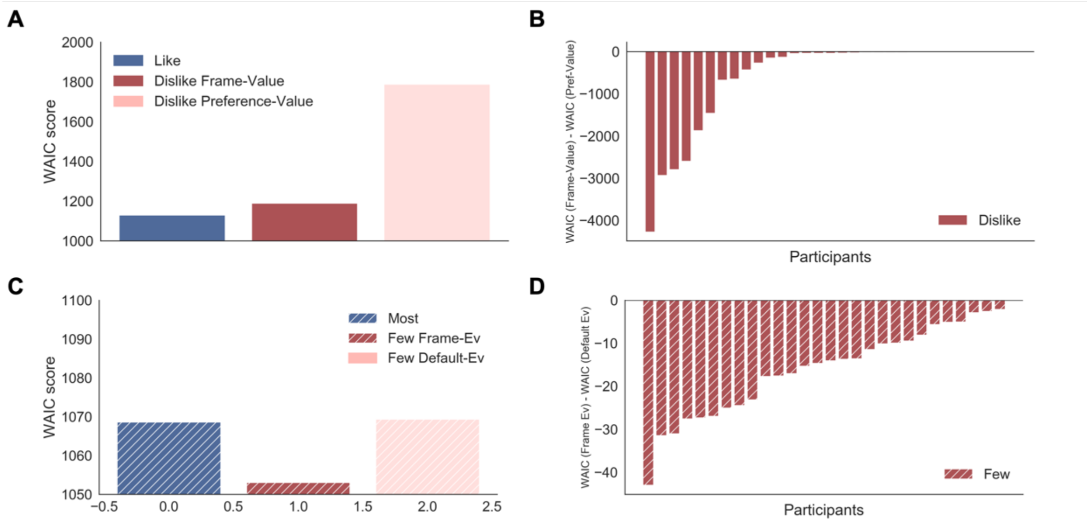


*Figure S14. GLAM model comparison. (A) Average WAIC scores for like and dislike GLAM models fitted at individual level. In the dislike frame, two possible models are compared: preference-value, value reported in the BDM bid was used directly to fit the data; and frame-value, value was adjusted to comply with the frame modification (see Methods for more details). The model accounting for goal-relevant evidence in the dislike frame had a better fit. (B) Individual WAIC differences between dislike models fitted with frame-value and preference-value. Negative differences indicate best fits for the frame-value in all the participants. (C) Average WAIC scores for most and fewest GLAM models fitted at individual level. In the fewest frame, two possible models are compared: default-evidence, the number of dots was used directly to fit the data, and frame-evidence, evidence was adjusted to comply with the frame modification (i.e., the opposite of the number of dots was used as evidence). (D) Individual WAIC differences between fewest models fitted with frame-evidence and default-evidence. Negative differences indicate best fits for the frame-evidence in all the participants. Solid colour indicates the value-based experiment and striped colours indicate the perceptual experiment.*

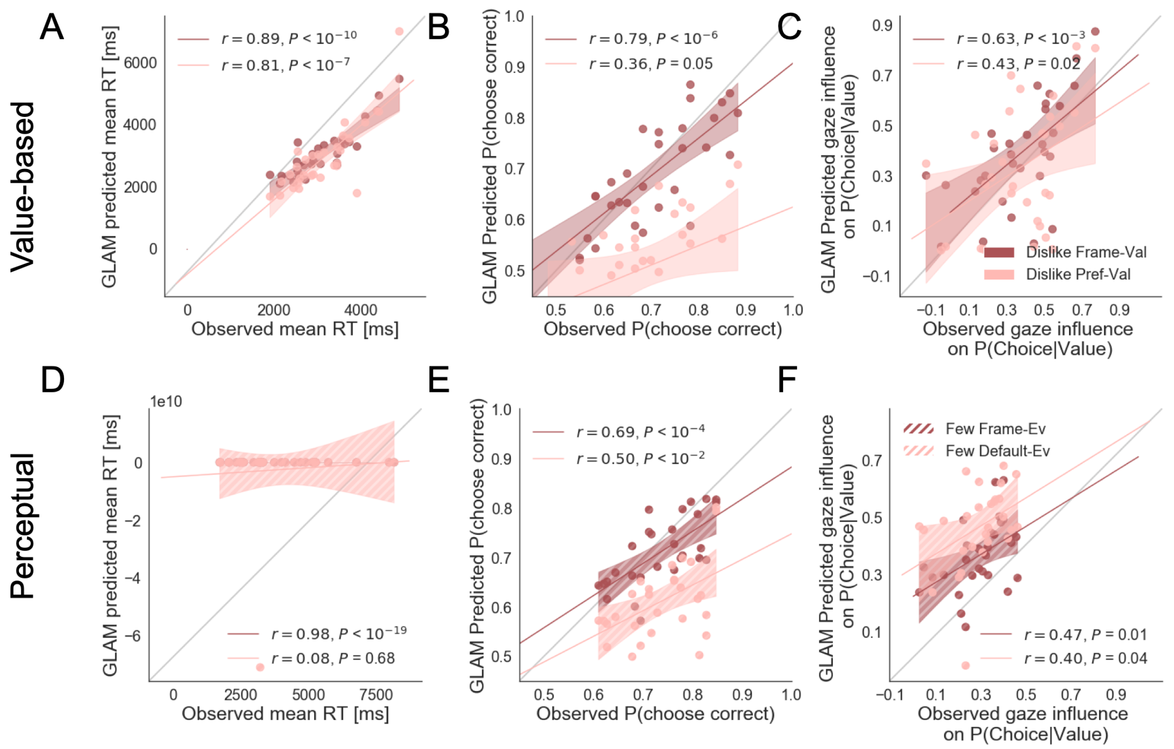


*Figure S15. Individual out-of-sample prediction from the GLAM model for behavioural measures in Value (dislike) (A-C) and Perceptual (fewest) Experiments (D-F). In the dislike frame, two models are used to generate simulations: preference-value, value reported in the BDM bid was used directly to fit GLAM model; and frame-value, the values were adjusted to comply with the frame modification. The model predicts participants mean reaction time (RT) (A), probability of choosing the best item (i.e., item with lower value) (B) and the influence of gaze in choice probability (C, check Results section for more details on gaze influence measure). The frame-value model correlates better with the observed data. In the Perceptual Experiment, fewest frame, also two possible models are used to generate simulations: default-evidence, the number of dots was used directly to fit the data, and frame-evidence, the evidence was adjusted to comply with the context modification (i.e., opposite of the number of dots). We show the correlation between the data and simulations for RT (D), the probability of choosing the best alternative (i.e., alternative with fewer dots) (E) and gaze influence (F). In this case, frame-evidence model also predicts the behaviour in the fewest frame better. The results corresponding to the model using frame-evidence are presented in red and the models using default-evidence in pink. Dots depict the average of predicted and observed measures for each participant. Lines depict the slope of the correlation between observations and the predictions. The shadowed region presents the 95% confidence intervals, with full colour representing Value Experiment and striped colour the Perceptual Experiment. Model predictions are simulated using parameters estimated from individual fits for even-numbered trials.*

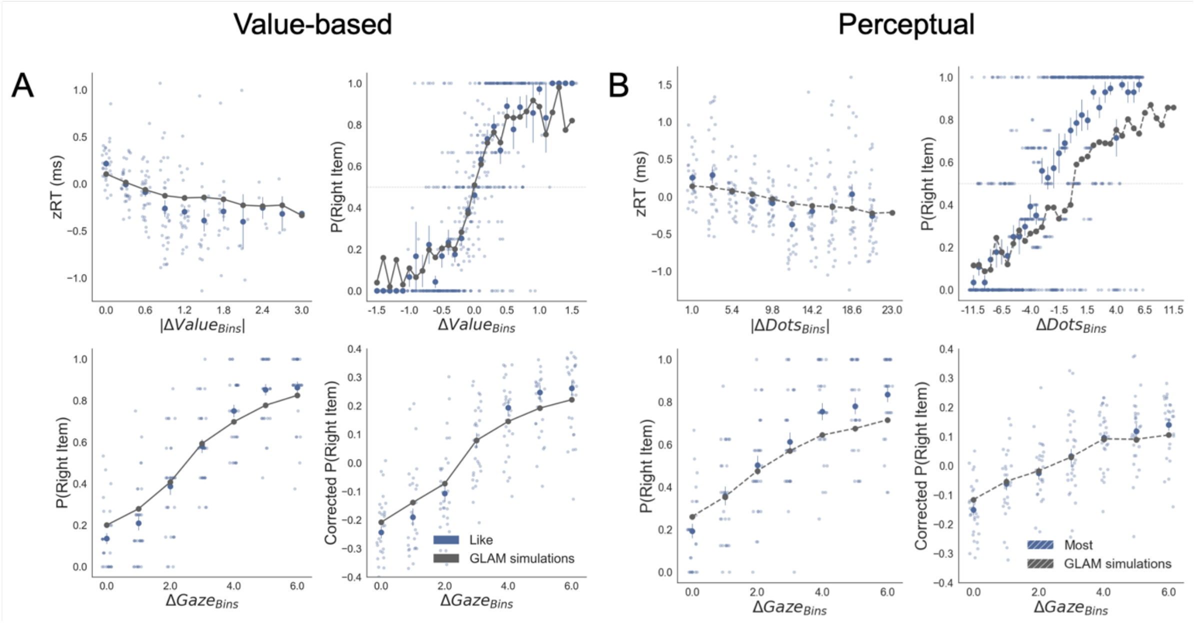


*Figure S16. Replication of behavioural effect of interest by simulations using the GLAM fitted for like (A) and most frames (B). The four panels present 4 relevant behavioural relationships found in the data: (top left) faster responses (shorter RT) when the choice is easier (i.e., easier choices are found with higher |ΔValue| in value-based and higher |ΔDots| in perceptual); (top right) probability of choosing the right alternative increases when the difference in evidence (value or number of dots) is higher in the alternative at the right side of the screen (ΔValue and ΔDots are calculated considering right minus left options); (bottom left) the probability of choosing an alternative depends on the gaze difference; and (bottom right) the gaze influence on choice depending on the difference in gaze time between both alternatives. Solid blue dots depict the mean of the data across participants in like and most frames. Light blue dots present the mean value for each participant. In the Value Experiment the solid grey lines show the average for model simulations. In the Perceptual Experiment segmented grey lines show the model simulations. Data is binned for visualization.*

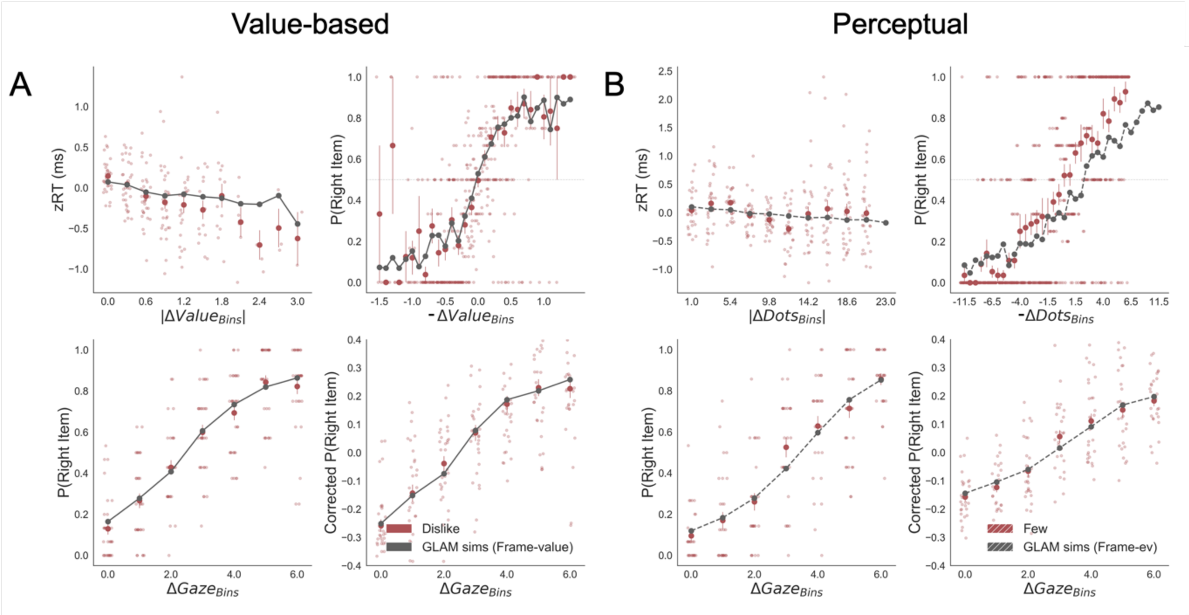


*Figure S17. Replication of behavioural effect of interest by simulations using the GLAM fitted for dislike (A) and fewest frames (B). Frame-relevant evidence was used to fit the model. The four panels present 4 relevant behavioural relationships found in the data. Top left: faster responses (shorter RT) when the choice is easier (i.e., easier choices are found with higher |ΔValue| in value-based and higher |ΔDots| in perceptual). Top right: probability of choosing the right alternative increases when the difference in evidence (value or number of dots) is lower in the alternative at the right side of the screen (notice that -ΔValue and -ΔDots are calculated considering left minus right options). Bottom left: the probability of choosing the right alternative depends on the gaze difference favouring the right option. Bottom right: the gaze influence on choice depending on the difference in gaze time between both alternatives. Solid red dots depict the mean of the data across participants in like and most frames. Light red dots present the mean value for each participant. In the Value Experiment the solid grey lines show the average for model simulations. In the Perceptual Experiment segmented grey lines show the model simulations. Data is binned for visualization*

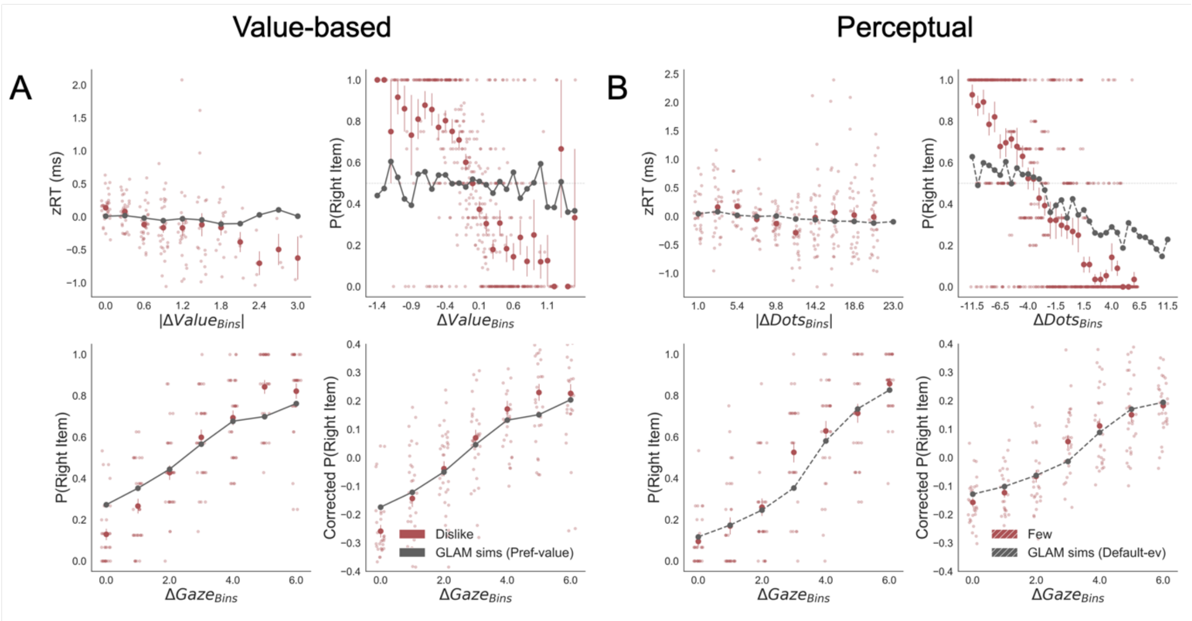


*Figure S18. Replication of behavioural effect of interest by simulations using the GLAM fitted for dislike (A) and fewest frames (B). In this case, the models were fitted without adapting the values and dot numbers to the evidence that was relevant for the particular frame i.e., the preference value and the default number of dots were used to fit the model in the dislike and fewest frame, respectively. The four panels present 4 relevant behavioural relationships found in the data. Top left: faster responses (shorter RT) when the choice is easier (i.e., easier choices are found with higher |ΔValue| in value-based and higher |ΔDots| in perceptual). Top right: probability of choosing the right alternative increases when the difference in evidence (value or number of dots) is lower in the alternative at the right side of the screen (ΔValue and ΔDots are calculated consider right minus left options). Bottom left: the probability of choosing the right alternative depends on the gaze difference favouring the right option. Bottom right: the gaze influence on choice depending on the difference in gaze time between both alternatives. No replication of the behavioural effect was found in this case for the relationship between RT -|ΔValue| and RT -|ΔValue| in dislike and fewest frames, respectively. Also P(right item)-ΔValue and P(right item)-ΔValue relationship was not replicated in dislike and fewest frames, respectively. Gaze effect seem to still keep its relationship, since gaze allocation time was not modified to account for the frame shift. Solid red dots depict the mean of the data across participants in like and most frames. Light red dots present the mean value for each participant. In the Value Experiment the solid grey lines show the average for model simulations. In the Perceptual Experiment segmented grey lines show the model simulations*. *Data is binned for visualization*

### SI 6: GLAM – Parameter Comparison

The results from the regression models presented in the *Results* section show that the nature of evidence integrated during the accumulation process depends on the frame in which participants make their choices. The Gaze-weighted Linear Accumulator Model (GLAM) predicts well participants' behaviour once frame-relevant evidence is employed to fit the model. Here we show the parameters obtained from this process. Four free parameters are fitted in GLAM: ν (drift term), γ (gaze bias), τ (evidence scaling) and σ (normally distributed noise standard deviation) [5]. For Value and Perceptual Experiments, we fitted the model in both frames and in each participant separately. The parameters were fitted using the even-numbered trials and in both studies the model fit was estimated using the WAIC score used to measure the fit of Bayesian Models (Figure S9).

*Value Experiment*. To explore variations in the process of accumulation of evidence characterized by GLAM, we compared the parameters obtained from the individual fit in *like* and *dislike* frames (Figure S14A). No significant variation between frames was found for the gaze bias (Mean $\gamma_{Like}$=-0.14, Mean $\gamma_{Dislike}$=0.03 , $\Delta\gamma_{Like-Dislike}$=-0.17, t=-1.66; p=0.11, ns), the scaling parameter ($\tau_{Like}$=2.81, $\tau_{Dislike}$=2.69, $\Delta\tau_{Like-Dislike}$=0.115 , t=0.313; p=0.75, ns) and the noise term (Mean $\sigma_{Like}$=0.0075, Mean $\sigma_{Dislike}$=0.0074, $\Delta\sigma_{Like-Dislike}$=0.00012, t=0.342; p=0.734, ns). We observed a significantly higher value of the drift term, ν, during the *like* frame ($\nu_{Like}=5.60\times10^{-5}$, $\nu_{Dislike}=4.53\times10^{-5}$, $\Delta\nu_{Like-Dislike}=1.06\times10^{-5}$, t=3.44; p<0.01). This means that evidence is accumulated faster during the *like* frame, which gives us an insight into the differences in the evidence accumulation product of the change frame modification.

*Perceptual Experiment*. We also compared the parameters obtained from GLAM individual fit in the perceptual experiment (Figure S14B). No significant variation between frames was found for the scaling parameter ($\tau_{Most}$=0.34, $\tau_{Few}$=0.13, $\Delta\tau_{Most-Few}$=0.212, t=1.43; p=0.16, ns) or the drift term (Mean $\nu_{Most}=3.8\times10^{-5}$, Mean $\nu_{Few}=3.99\times10^{-5}$, $\Delta\nu_{Most-Few}=-1.92\times10^{-6}$, t=-0.465; p=0.645, ns). The gaze bias is larger during the *fewest* frame ($\gamma_{Most}$=0.48, $\gamma_{Few}$=0.26, $\Delta\gamma_{Most-Few}$=0.22, t=2.61; p<0.05). The σ parameter is also significantly different depending on the frame, with higher noise in the *most* frame ($\sigma_{Most}$=0.0073, $\sigma_{Few}$=0.0066, $\Delta\sigma_{Most-Few}$=0.0007, t=2.26; p<0.05). In summary, the accumulation process seems to be noisier and less affected by visual attention in the *most* frame. In both frames, the finding that γ<1 indicates that gaze modulates the accumulation of evidence.

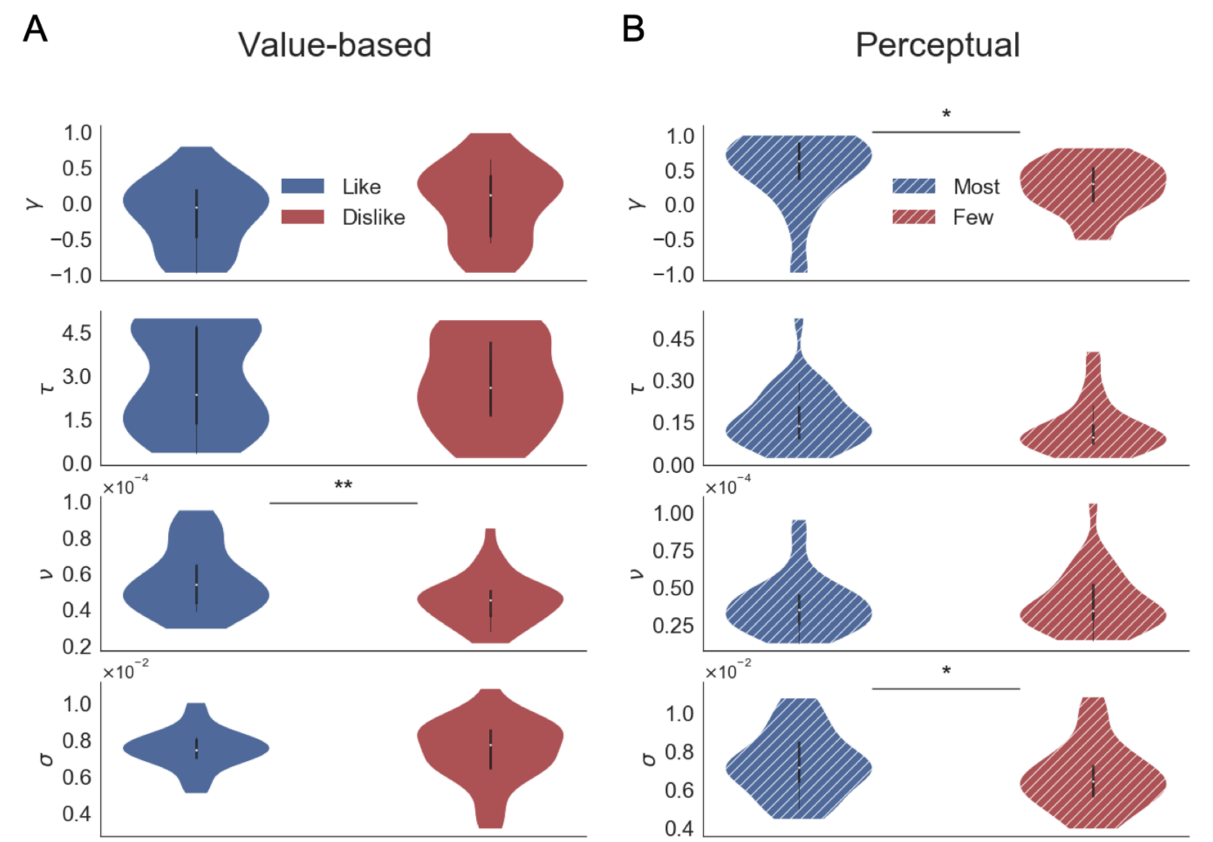


*Figure S19. Parameters fitted at subject level using GLAM in Value (A) and Perceptual (B) Experiments. The free parameters are γ (gaze bias), τ (evidence scaling), v (drift term) and σ (standard deviation of the normally distributed noise). In the Value Experiment we found a significant decrease in the drift term during the dislike frame, maybe indicating a more uncertain decision process. The parameters in Perceptual Experiment were significantly different for gaze bias and noise term, with higher γ and σ values in the most frame. This may indicate a reduced effect of gaze on choice during the most frame and slightly less noisier accumulation process in the fewest frame. In each experiment, the GLAM parameters were fitted independently for each frame. In the violin plot, red and blue areas indicate the distribution of the parameters across participants. Black bars present the 25, 50 and 75 percentiles of the data. Solid colour indicates the Value Experiment and striped colours indicate the Perceptual Experiment.*

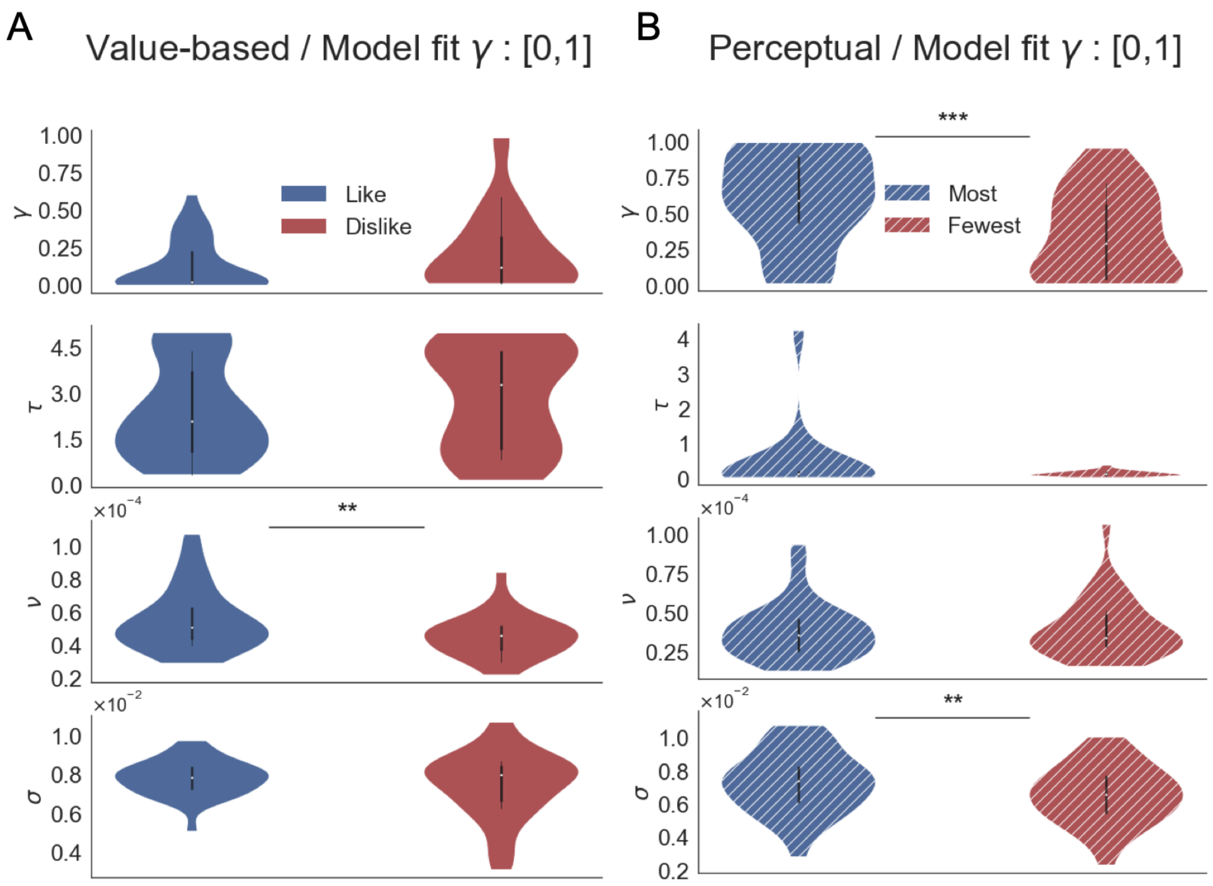


*Figure S20. GLAM model parameters when the model fit is performed constraining γ to [0,1] range. Thomas et al. [5] describes a "leakage" of evidence when γ<0, which can be a conflicting assumption in this type of models. We corroborated that the differences between the parameters in like/dislike and most/fewest remain the same in comparison to the fit reported constraining γ to [-1,1].*

### SI 7: Attentional Drift Diffusion Model

The attentional Drift Diffusion Model (aDDM) has been extensively used in literature to characterise the effect of attention over choice [4]. Unlike GLAM, aDDM considers the dynamics of fixations during trials to fit the model. To further support our idea that goal-relevant evidence is accumulated, we fitted both Value and Perceptual datasets with the aDDM model, as implemented by Tavares et al. [6] (aDDM toolbox, https://github.com/goptavares/aDDM-Toolbox).

The aDDM model assumes that evidence is accumulated dynamically in a variable called the relative decision value (RDV) signal. RDV starts at 0 and it evolves over time, accumulating evidence until a barrier is reached (+1 or -1) which will define the alternative to be selected (right or left). Every time step, RDV changes according to $\mu\Delta t + \varepsilon t$, with μ the deterministic change (slope term) and ε the Gaussian noise term. The fixation to the two alternatives will define the value of μ: when the left option is fixated $\mu = d(r_{left} - \theta r_{right})$ and $\mu = d(r_{right} - \theta r_{left})$ for the right option. Therefore, the aDDM model considers three free parameters: d, σ and $\theta$. The parameter d is a positive constant characterising the speed of integration; σ is the standard deviation for a zero-mean Gaussian distribution for noise, and θ is the attentional parameter that controls the size of the attentional bias (range between 0 to 1). If θ=1, the model is reduced to a standard drift-diffusion model (DDM) without attentional bias.

*Group model fitting*. The models were fitted to choice and RT data independently for *like* and *dislike* frames in our Value Experiment and for *most* and *fewest* frames in the Perceptual Experiment. The odd trials of the pooled data from 31 participants in value-based data and 32 participants for perceptual case was used to fit the models. The model considers the available evidence (item value and number of dots) and the sequence of fixations for each trial. As in GLAM, we fitted the parameters in *dislike* and *fewest* frames considering a version of the input values/evidence that accounted for the change in the objective of the task (i.e., reporting item not preferred or the alternatives with fewer dots, respectively). To compare, we also fitted another model using the evidence "by default" (i.e., BDM bid values or number of dots in the circles). To account for the different ranges of item valuation used by the participants we normalized the value reports by binning the item values at a participant level. In the Value Experiment, the data were separated in 6 bins using quantiles-based discretization. In the perceptual case, given the distribution of the evidence (i.e three numerosity levels and smaller dots differences between two alternatives) we separated the dots data in 8 bins. The maximum likelihood estimation (MLE) procedure was carried in iterative steps searching over a grid with the 3 model parameters. Initial grid was set to [0.001, 0.005, 0.01] for d, [0.01, 0.05, 0.1] for σ and [0.01, 0.5, 1] for θ. The likelihood for choice and RT in odd-trials, conditional to the pattern of fixations observed in that trial, was calculated for each combination of parameters in the grid (check Tavares et al. [6] for the details of the algorithm to simulate aDDM trials). The time step used for the estimation of aDDM was 10 ms. The set of parameters with lower negative log-likelihood (NLL) was used as center of the grid for the next iteration. Therefore, the grid to search in the next iteration (t+1) was defined as $[d_t - \Delta d_t/2, d_t, d_t + \Delta d_t/2]$, $[\theta_t - \Delta\theta_t/2, \theta_t, \Delta\theta_t/2]$, and $[\sigma_t - \Delta\sigma_t/2, \sigma_t, \sigma_t + \Delta\sigma_t/2]$, considering the respective constrains of each parameter value. The iterative process finished once the improvement in the MLE of the proposed parameter solution was smaller than 0.05% ($|minNNL_{t+1} - minNNL_t| < 0.0005 * minNNL_t$). The convergence was reached after two iterations in our models. In our results, we found that for both, *dislike* and *fewest* conditions, the model fitted using goal-relevant evidence had better performance than the model using default estimated value or number of dots, as indicated by a lower NLL value.

*Table S7. aDDM model parameters. Estimated parameters for Value and Perceptual Experiments. Parameter description - d: speed of integration; σ: standard deviation for the noise distribution, θ :attentional bias. NNL: negative log-likelihood of the models indicating goodness-of-fit.*

| | Value-based | | | Perceptual | | |
|---|---|---|---|---|---|---|
| | Like | Dislike Preference-values | Dislike Frame-values | More | Fewest Default-evidence | Fewest Frame-evidence |
| d | 0.001 | 0 | 0.001 | 0.001 | 0.001 | 0.001 |
| σ | 0.05 | 0.05 | 0.05 | 0.05 | 0.05 | 0.05 |
| *θ* | 0 | 0 | 0 | 0.255 | 0 | 0.01 |
| NLL | 12441.012* | 13342.297 | 12640.837* | 13948.411* | 14169.154 | 13826.983* |

* Indicates the model with lower NLL for that frame

*Out-of-sample group simulations.* To test the capacity of the model to predict out-of- sample, the aDDM with the best fitted parameters using odd-numbered trials was used to predict the behaviour observed on the even-numbered trials. We did 40000 simulations for the Value Experiment and 48000 trials for the Perceptual Experiment. Fixations, latencies and inter-fixations transitions were sampled from empirical distributions, obtained from the pooled even-numbered trials across participants following the procedure used by Tavares and colleagues [6].

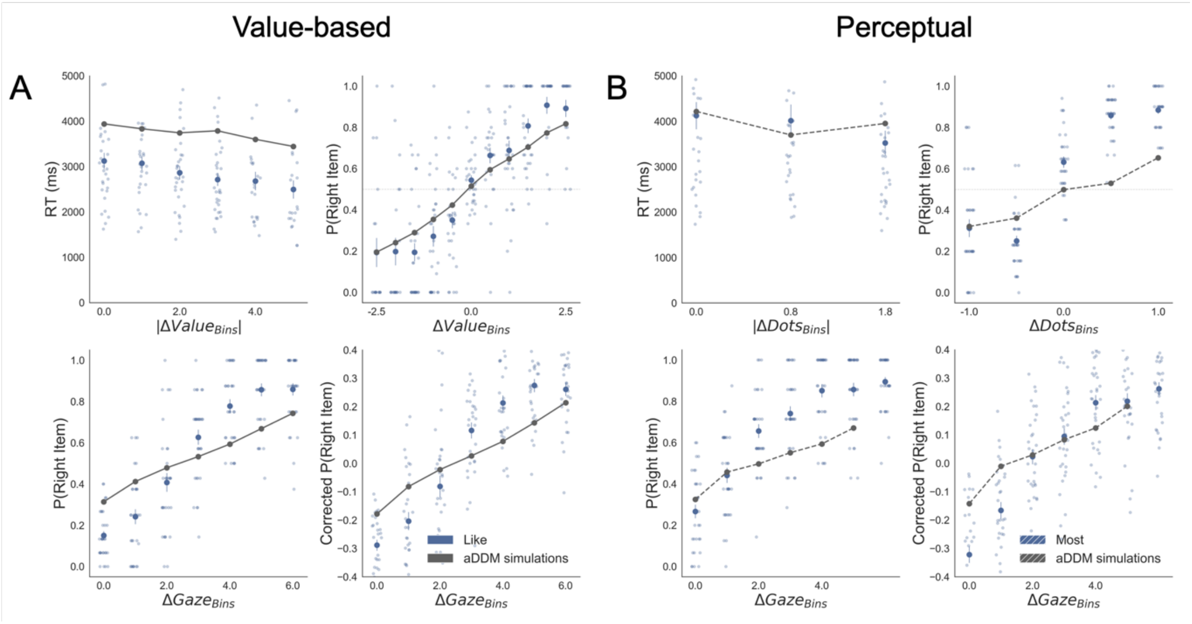


*Figure S21. Replication of behavioural effects by aDDM simulations for like (A) and most frames (B). The four panels present 4 relevant behavioural relationships found in the data. Top left: faster responses (shorter reaction time, RT) when the choice is easier (i.e., easier choices are found with higher |ΔValue| and |ΔDots| in Value an Perceptual Experiments, respectively). Top right: probability of choosing the right alternative increases when the evidence towards the right item is higher (ΔValue and ΔDots are calculated considering right minus left options). Bottom left: the probability of choosing the item on the right side of the screen depends on the gaze time difference (ΔGaze, calculated as the time observing the right minus the left item). Bottom right: gaze influence on choice depending on the difference in ΔGaze (check Results section for more details on gaze influence). Solid blue dots depict the mean of the data across participants in like and most frames. Light blue dots show the mean value for each participant. In Value Experiment the solid grey lines show the average for model simulations. In the Perceptual Experiment segmented grey lines show the average for model simulations. Data and simulations were binned for visualization.*

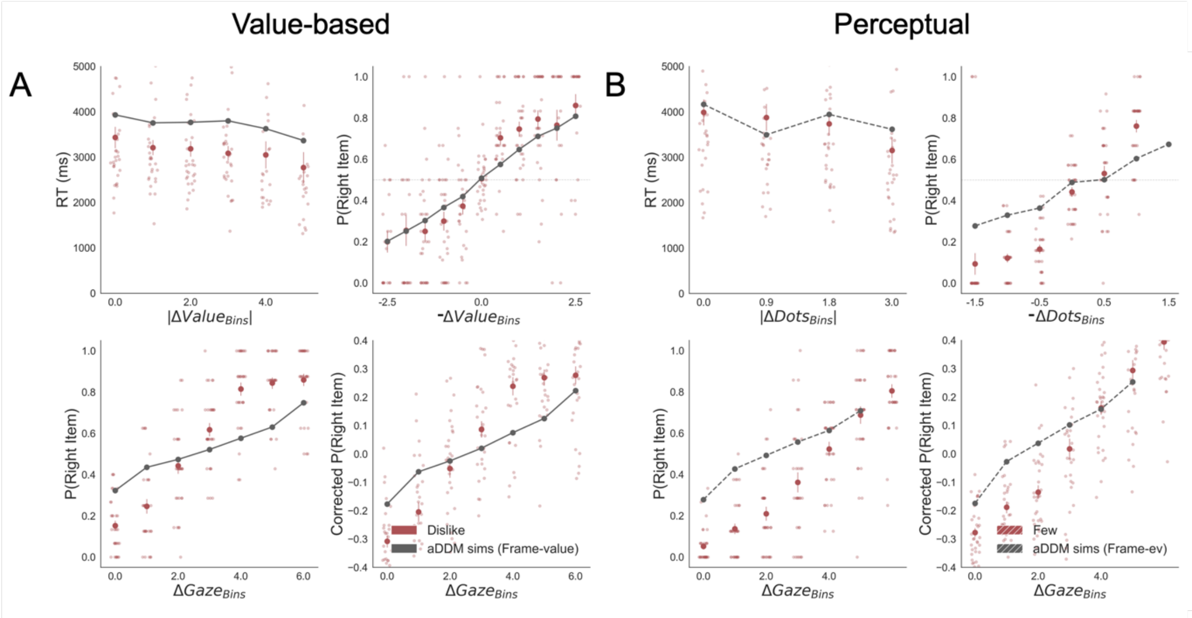


*Figure S22. Replication of behavioural effects by aDDM simulations for dislike (A) and fewest (B) frames. Importantly, these models were fitted using goal-relevant evidence. The four panels present 4 relevant behavioural relationships found in the data. Top left: faster responses (shorter reaction time, RT) when the choice is easier (i.e., easier choices are found with higher |ΔValue| and |ΔDots| in Value and Perceptual Experiments, respectively). Top right: probability of choosing the right alternative increases when the evidence towards the left item is higher (-ΔValue and –ΔDots, i.e., increment when left item is more valuable or has more dots than the right item). Bottom left: the probability of choosing the item on the right side of the screen depends on the gaze time difference (ΔGaze, calculated as the time observing the right minus the left item). Bottom right: gaze influence on choice depending on the difference in ΔGaze (check Results section for more details on gaze influence). Solid red dots depict the mean of the data across participants in like and most frames. Light red dots show the mean value for each participant. In Value Experiment the solid grey lines show the average for model simulations. In the Perceptual Experiment segmented grey lines show the average for model simulations. Data and simulations were binned for visualization.*

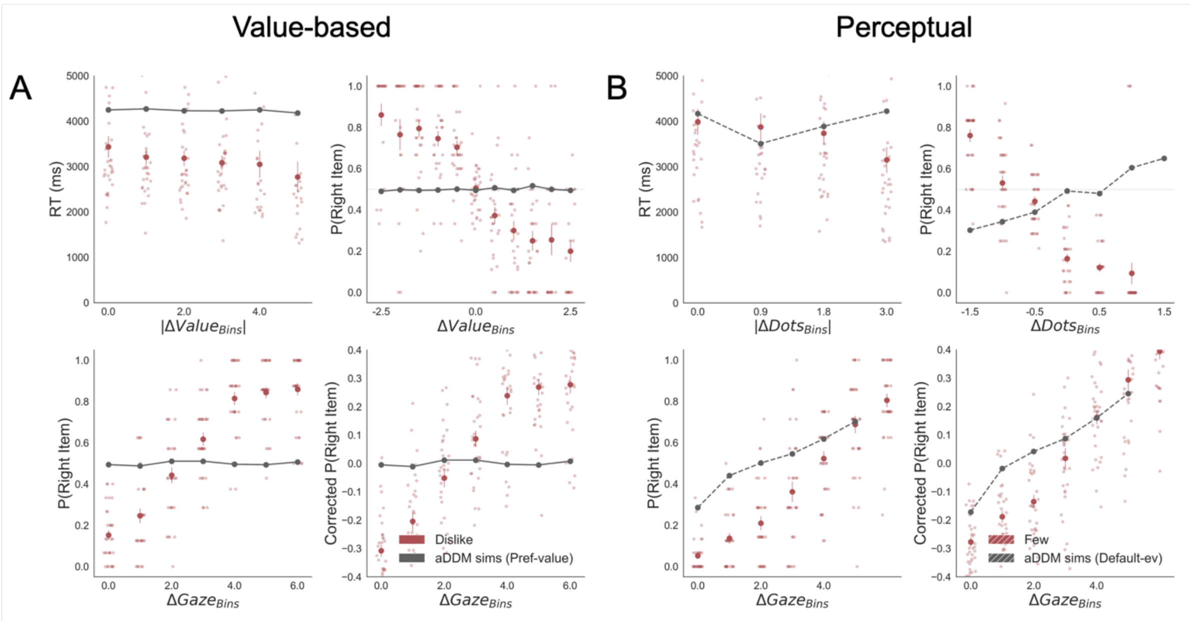


*Figure S23. Replication of behavioural effects by aDDM simulations for dislike (A) and fewest frames (B). Importantly, these models were fitted using the default evidence in Value and Perceptual Experiments, i.e., preference value and number of dots, respectively. Unlike the models fitted with goal-relevant evidence, these models do not capture reaction time (RT) and choice behaviour in dislike and fewest frames. The four panels present 4 relevant behavioural relationships found in the data. Top left: faster responses (shorter RT) when the choice is easier (i.e., easier choices are found with higher |ΔValue| and |ΔDots| in Value an Perceptual Experiments, respectively). Top right: probability of choosing the right alternative increases when the evidence towards the left item is higher (ΔValue and ΔDots are calculated considering right minus left options). Bottom left: the probability of choosing the item on the right side of the screen depends on the gaze time difference (ΔGaze, calculated as the time observing the right minus the left item). Bottom right: gaze influence on choice depending on the difference in ΔGaze (check Results section for more details on gaze influence). Solid blue dots depict the mean of the data across participants in like and most frames. Light blue dots show the mean value for each participant. In Value Experiment the solid grey lines show the average for model simulations. In the Perceptual Experiment segmented grey lines show the average for model simulations. Data and simulations were binned for visualization.*

### SI 8: GLAM – Balance of Evidence Simulations

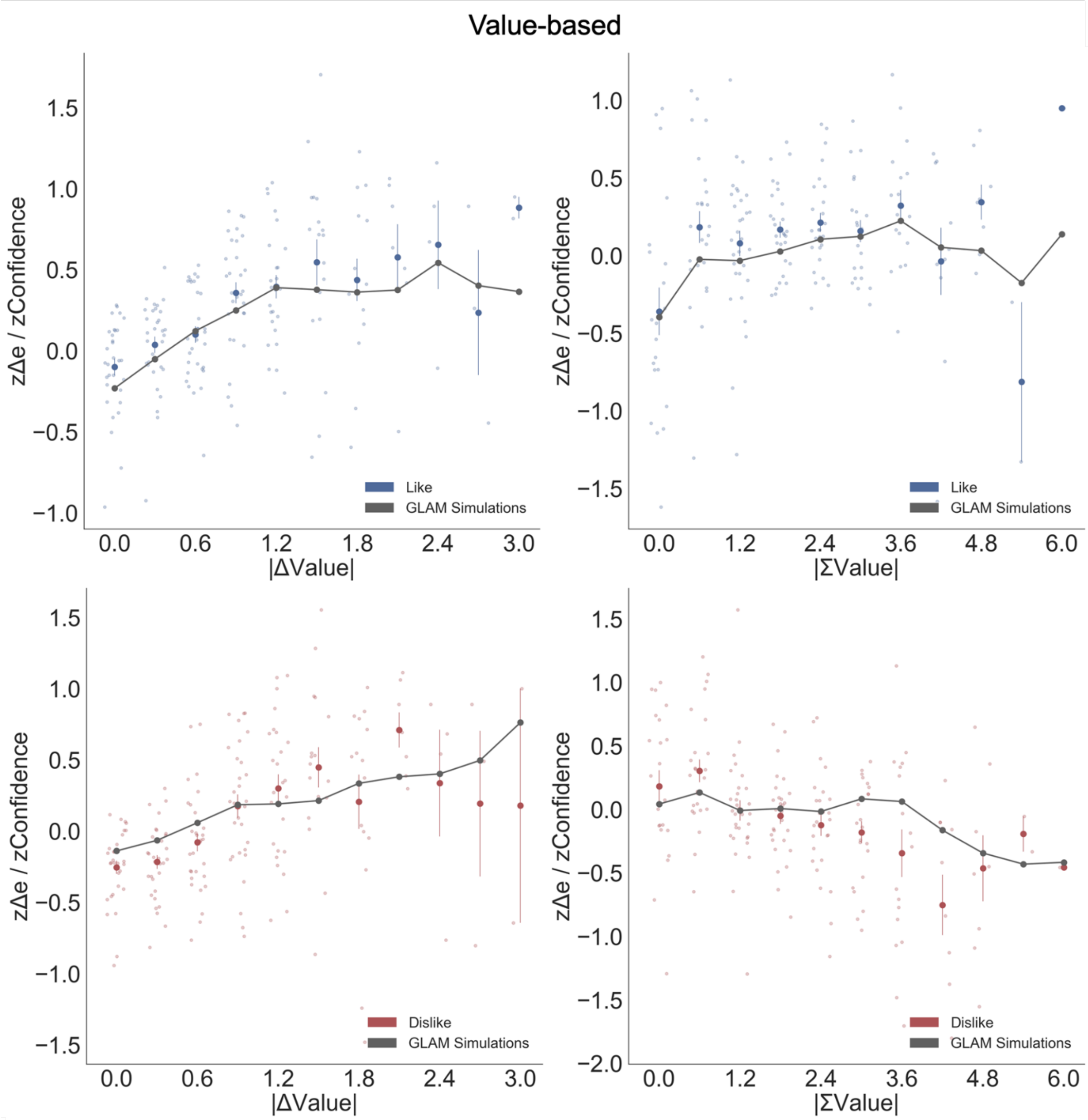


*Figure S24. Balance of evidence simulations in the Value Experiment. The difference between accumulators (Δe) obtained from GLAM simulations matches participants' confidence. Top left: a higher value difference between the two items (|ΔValue|) increases confidence and simulated Δe. Top right: in the like frame, an increase in the summed value of the two alternatives (|ΣValue|) boosts confidence and simulated Δe. Bottom left: as in like frame, |ΔValue| boosted confidence and Δe in dislike frame. Bottom right: in the dislike frame, the effect of |ΣValue| over confidence flips: confidence and Δe decrease with higher values of the alternatives, accounting for the change in goal. Blue and red dots depict the (z-scored) confidence taken from participants in like and dislike frames (respectively). Grey line presents the model simulations for both separate frames. Data was segmented in 11 bins for ΔValue or ΣValue.*

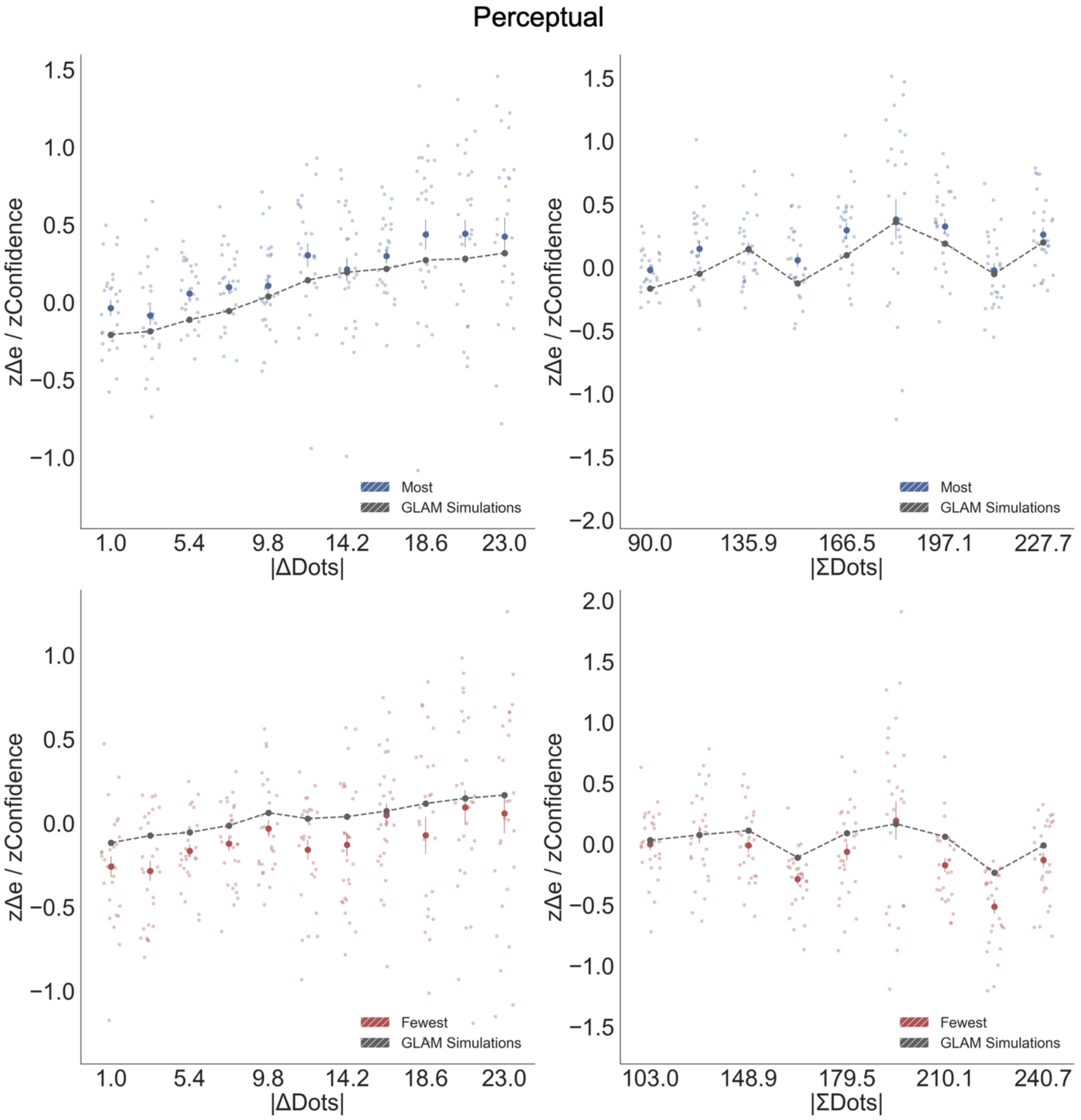


*Figure S25. Balance of evidence simulations in the Perceptual Experiment. As in Value Experiment, the difference between accumulators (Δe) obtained from GLAM simulations matches participants' confidence. Top left: a higher difference in number of dots between the two circles (|ΔDots|) increases confidence and simulated Δe. Top right: in the most frame, an increase in the summed number of dots (|ΣDots|) boosts confidence and simulated Δe. Bottom left: as in most frame, |ΔDots| boosted confidence and Δe in fewest frame. Bottom right: in the fewest frame, the effect of |ΣDots| over confidence flips: confidence and Δe decrease with higher number of dots in both circles, accounting for the change in goal. Blue and red dots depict the (z-scored) confidence taken from participants in like and dislike frames (respectively). Grey line presents the model simulations for both separate frames. Data was segmented in 11 bins for ΔValue or ΣValue.*

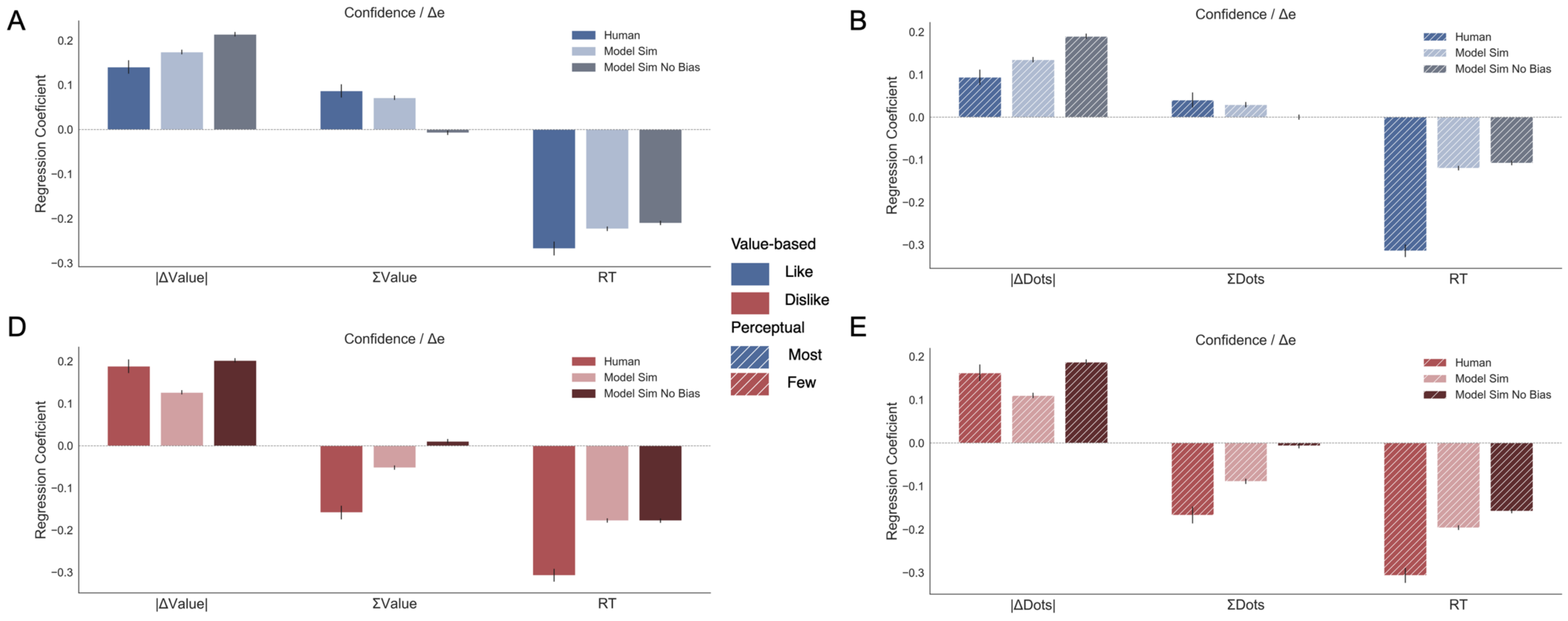


*Figure S26. Pooled linear regressions to predict balance of evidence (Δe) simulations. Here the full model results for figure 6 (see Results section) are displayed. In Value Experiment, the full simulations of Δe replicated the pattern of results obtained in human data (confidence results), i.e., there is a flip in the sign of ΣValue effect over confidence between like (A) and dislike (D) frames. However, if the gaze asymmetry is removed we found the effect of ΣValue over Δe disappears. The results in Perceptual Experiment, most (B) and fewest (E) frames, mirror the findings in Value Experiment.*

### SI 9: Normative Model – Proof of Propositions 1 and 2

We begin with the proof of Proposition 1.

Recall that agents start with an identical, Normal prior over the quality $v_i$ of each object; and that they receive signals $x_i = v_i + \epsilon_i$, with $\epsilon_i$ independently and identically distributed with $\epsilon_i \sim N(0, \sigma_\epsilon^2)$. This implies that, after receiving the first signal, the belief about $v_i$ is again normally distributed: denote $\mu_i$ and $\sigma_v^2$ the mean and variance of these posterior beliefs. Note that $\sigma_v^2$ is the same for all $i$.

After the agent acquires a second signal about item $i$, this generate a posterior about $v_i$; denote by $\mu_{i,2}$ the mean of such posterior. Note that, after the first signal but before acquiring a second signal, the agent's belief about the distribution of $\mu_{i,2}$ is $N(\mu_i, \theta)$, for some $\theta > 0$ independent of $i$. Note also that, by construction, for $i \neq i_1$ we have

$$V(i) = \max\{\mu_{i_1}, \mu_{i,2}\},$$

while for $i_1$ we have

$$V(i_1) = \max\{\mu_{i_2}, \mu_{i_1,2}\}.$$

Denote by $F_i$ the agent's corresponding belief on the distribution of $V(i)$. For $i \neq i_1$, $F_i$ coincides with $N(\mu_i, \theta)$ for values above $\mu_{i_1}$, but has a mass point at $\mu_{i_1}$ equal to the probability that $N(\mu_i, \theta)$ is below $\mu_{i_1}$. That is, for any Borel subset $A$, if $A \subseteq (\mu_{i_1}, +\infty)$, then $F_i(A) = N(\mu_i, \theta)(A)$; [1] while $F_i(\mu_{i_1}) = N(\mu_i, \theta)([-\infty, \mu_{i_1}])$.

Similarly, $F_{i_1}$ coincides with $N(\mu_{i_1}, \theta)$ for values above $\mu_{i_2}$, but has a mass point at $\mu_{i_2}$ equal to the probability that $N(\mu_i, \theta)$ is below $\mu_{i_2}$. That is, for any Borel $A \subset \mathbb{R}$, if $A \subseteq (\mu_{i_2}, +\infty)$, then $F_{i_1}(A) = N(\mu_{i_1}, \theta)(A)$; while $F_{i_1}(\mu_{i_2}) = N(\mu_{i_1}, \theta)([-\infty, \mu_{i_2}])$.

*Claim 1.* $\mathbb{E}[V(i_1)] = \mathbb{E}[V(i_2)]$.

*Proof.* Following our notation above and denoting $f_\mu$ the Probability Density function of $N(\mu, \theta)$, we have

$$\mathbb{E}[V(i_1)] = \int_{-\infty}^{+\infty} x\, dF_{i_1}(x) = \int_{-\infty}^{\mu_{i_2}} \mu_{i_2} f_{\mu_{i_1}}(x)dx + \int_{\mu_{i_2}}^{+\infty} x\, f_{\mu_{i_1}}(x)dx,$$

---

[1] We use $N(\mu_i, \theta)$ to denote the probability measure, thus $N(\mu_i, \theta)(A)$ indicates the probability of $A$.

$$\mathbb{E}[V(i_2)] = \int_{-\infty}^{+\infty} x\, dF_{i_2}(x) = \int_{-\infty}^{\mu_{i_1}} \mu_{i_1} f_{\mu_{i_2}}(x)dx + \int_{\mu_{i_1}}^{+\infty} x\, f_{\mu_{i_2}}(x)dx.$$

By construction we have

$$\mu_{i_1} = \int_{-\infty}^{\mu_{i_2}} x\, f_{\mu_{i_1}}(x)dx + \int_{\mu_{i_2}}^{+\infty} x\, f_{\mu_{i_1}}(x)dx.$$

It follows that

$$\mathbb{E}[V(i_1)] - \mu_{i_1} = \int_{-\infty}^{\mu_{i_2}} (\mu_{i_2} - x)\, f_{\mu_{i_1}}(x)dx \qquad \text{(Eq. 1)}$$

and

$$\mathbb{E}[V(i_2)] - \mu_{i_1} = \int_{\mu_{i_1}}^{+\infty} \left(x - \mu_{i_1}\right) f_{\mu_{i_2}}(x)dx. \qquad \text{(Eq. 2)}$$

But note that

$$\int_{-\infty}^{\mu_{i_2}} (\mu_{i_2} - x)\, f_{\mu_{i_1}}(x)dx = \int_{2\mu_{i_1}-\mu_{i_2}}^{+\infty} (x - 2\mu_{i_1} + \mu_{i_2})\, f_{\mu_{i_1}}(x)dx = \int_{\mu_{i_1}}^{+\infty} (x - \mu_{i_1})\, f_{\mu_{i_2}}(x)dx.$$

With Eq. 1 and 2, this proves the claim. ■

*Claim 2*. If N>2, $\mathbb{E}[V(i_2)] > \mathbb{E}\left[V\left(i_j\right)\right]$ for all $j > 2$.

*Proof*. To prove the claim we show that $F_{i_2}$ First Order Stochastically Dominate $F_{i_j}$ for all $j > 2$. Let $\delta =: \mu_{i_2} - \mu_{i_j}$ and, for any Borel $A \subset \mathbb{R}$, denote $A + \delta \coloneqq \{x + \delta : x \in A\}$. Note that we have $\delta > 0$ and that $N\left(\mu_{i_2}, \theta\right)(A + \delta) = N\left(\mu_{i_j}, \theta\right)(A)$. It follows that for all $A \subseteq \left(\mu_{i_1}, +\infty\right)$, $F_{i_j}(A) = F_{i_2}(A + \delta)$. That is: above $\mu_{i_1}$, $F_{i_2}$ assigns the same weight to strictly higher values. Both distributions are also bounded below by $\mu_{i_1}$. Moreover,

$$F_{i_2}\left(\mu_{i_1}\right) = N\left(\mu_{i_2}, \theta\right)\left([-\infty, \mu_{i_1}]\right) < N\left(\mu_{i_j}, \theta\right)\left([-\infty, \mu_{i_1}]\right) = F_{i_j}\left(\mu_{i_1}\right).$$

It follows that $F_{i_2}$ First Order Stochastically Dominate $F_{i_j}$ for all $j > 2$. The claim then follows. ■

The two claims together prove Proposition 1.

The proof of Proposition 2 is very similar once we replace $i_j$ by $i_{N+1-j}$ for $j = 1, \ldots, N$: the problem of maximizing the expected utility of the remaining items is strategically equivalent to the problem of choosing the lowest item, which, in turn, is symmetric to the problem of choosing the best item. $QED$.
